# Beyond Fixation: Persistent Genetic Variation Under Intense Selection

**DOI:** 10.64898/2026.03.02.706684

**Authors:** Kenneth R. Arnold, Zachary S. Greenspan, Ryan D. Robinson, Anastasia Pupo, Valeria V. Chavarin, Kevin S. Chang, Corrina O. Cannell, Miao Qi, Laurence D. Mueller, Michael R. Rose, Mark A. Phillips

## Abstract

Understanding how and why genetic variation is maintained under sustained selection remains a central question in evolutionary genetics. Experimental evolution shows that adaptation in sexually reproducing populations is often highly polygenic, proceeding through coordinated, genome-wide allele frequency shifts from standing variation rather than classic hard sweeps. Recent explanations emphasize highly polygenic architectures, optimizing-selection, and genetic redundancy, which can slow fixation by distributing selection across many loci during adaptation. However, observations from long-term selection experiments reveal a pattern these frameworks do not fully explain: substantial genetic variation persists after hundreds of generations of intense directional selection in constant environments. Here, we use long-term experimental evolution in *Drosophila melanogaster* to test whether balancing-selection actively maintains genetic variation under strong life-history selection and preserves evolutionary reversibility. Longstanding populations selected for accelerated or delayed reproduction were shifted to the opposing regime, imposing age-structured fitness trade-offs. Notably, selection for early reproduction is associated with substantial loss of genetic variation, providing a stringent test of whether standing variation is truly depleted. Following reciprocal shifts, populations showed rapid phenotypic convergence toward the target regime. At the genomic level, allele-frequency trajectories were strongly antiparallel and highly repeatable across replicates, revealing coordinated polygenic responses. Relaxing long-standing early-life selection produced a pronounced rebound in genome-wide heterozygosity. Deep sequencing uncovered ultra-rare alleles at sites appearing fixed under standard coverage, indicating low-frequency functional variation persists below detection thresholds. These results suggest that substantial genetic variation can persist under intense directional selection and be rapidly redeployed when selection reverses, consistent with widespread balancing-selection.

## Introduction

Understanding how genetic variation is shaped and maintained in populations has long been a central question within evolutionary genetics (Lewontin 1974). A major puzzle is why substantial genetic variation often persists within sexually reproducing populations despite long-term, intense selection, under which classical models predict rapid fixation and loss of diversity. Evolution experiments in sexually reproducing organisms have repeatedly shown that responses to selection are often highly polygenic, fueled by standing genetic variation, and characterized by modest coordinated shifts in allele frequencies rather than classic hard sweeps in which selected alleles and linked sites fix (Burke 2012; Barghi and Schlötterer 2020). In mesocosm experiments mimicking natural conditions, the absence of sweeps and persistence of genetic variation is often attributed to fluctuating selection and adaptive tracking (Rudman et al. 2022; Bitter et al. 2024). However, we know that this explanation cannot be applied to laboratory-based experimental evolution studies where environments and selection pressures are typically constant.

Experimental evolution combined with whole-genome sequencing (“E&R”) is an established approach for studying the genetic basis of adaptation and complex trait variation (Long et al. 2015; Schlötterer et al. 2015). Across the *Drosophila* studies that comprise most of this work, the shift versus sweep dynamic is widespread (e.g. Burke et al. 2010; Tobler et al. 2014; Graves et al. 2017; Barghi et al. 2019, 2020). It is even observed in outcrossing yeast studies, where large population sizes and longer selection durations allow for a greater possibility of adaptation from de novo mutations (Burke et al. 2014; Phillips et al. 2020; Linder et al. 2022). Recent explanations for the “shifts not sweeps” pattern emphasize highly polygenic architectures, optimizing selection, and genetic redundancy (Chevin and Hospital 2008; Barghi et al. 2019; Hayward and Sella 2022). However, while this framework is compelling, it does not resolve a puzzling pattern that emerges in the case of long-term selection experiments, where populations experience sustained selection for many hundreds of generations. Although genetic redundancy can slow fixation by distributing selection across many loci during ongoing adaptation, it does not provide a mechanism for actively maintaining polymorphisms once populations approach a phenotypic optimum, at which point drift is expected to erode variation.

Evidence for this phenomenon comes from our own long-term *Drosophila* experimental evolution system, in which populations maintained under intense selection for over 1,000 generations retain substantially more genetic variation than expected under neutral drift and under forward simulations incorporating both directional and optimizing selection (Phillips et al. 2016). Why so much genetic variation persists after hundreds of generations of intense selection remains an open question. Here we propose that pervasive antagonistic pleiotropy may play an important role in maintaining genetic variation by constraining fixation and slowing the erosion of polymorphism under sustained selection.

In previous work, we raised the possibility that the long-term persistence of genetic variation in these populations reflects pervasive balancing selection acting within populations rather than passive retention (Graves et al. 2017). This idea was motivated by a key result from that study: despite many generations of selection on reproductive timing, populations recently derived from the same ancestral stock rapidly converged phenotypically and genotypically with long-standing selected lines. This repeatability implies that adaptive alleles are retained in the founder populations far longer than expected, given that hundreds of generations of selection and drift should otherwise erode standing variation. Antagonistic pleiotropy provides a plausible mechanism: alleles that enhance fitness through one life-history character may impose costs upon other characters, constraining fixation, and in some cases, sustaining genetic polymorphism over time (Rose 1982; Rose 1985; Brud and Guerrero 2026).

Here we aim to empirically test whether widespread balancing selection contributes to the long-term maintenance of genetic variation in *Drosophila* experimental evolution. We focus on populations that have been under intense selection for early or late reproduction for hundreds of generations and assess whether they retain the capacity to evolve back toward ancestral phenotypic and genomic states when selection is reversed.

The early-reproduction populations, termed the A-types, are maintained on a 10-day generation cycle, whereas the late-reproductive populations, the C-types, are maintained on a 28-day cycle (Table S1). Both sets were ultimately descended from the same ancestral Ives base population (Ives 1970; Rose et al. 2004). A-type selection produces accelerated development, reduced adult lifespan, reduced stress resistance, increased early-life fecundity, and distinct metabolic profiles (Burke et al. 2016; Kezos et al. 2023; Hubert et al. 2025). Genomic characterization further indicates that these phenotypes are underlain by widespread genomic responses to selection, including the apparent fixation of some polymorphic sites and regions of the genome where variation is almost entirely purged. (At present, the longest-standing A-type populations have experienced ∼1,200 generations of selection for early reproduction, and the more recently derived A-types ∼650). Given that population sizes are on the order of a thousand, beneficial *de novo* mutations are unlikely to be a factor driving adaptation and populations should have reached mutation-selection-drift equilibrium under their respective selection regimes. Under standard expectations when antagonistic pleiotropy is absent, most segregating variation should be selectively neutral or weakly deleterious, slowly drifting toward fixation or loss. Thus, the ability of A-type populations to move rapidly back toward C-type states when selection is altered would imply that genetic variation has been actively maintained at equilibrium frequencies, rather than “passively” retained during incomplete selection.

To assess the possible conclusion described above, we use antiparallel selection shifts between the A- and C-type regimes and follow two trajectories: A→C (A-types placed under C-type conditions) and C→A (C-types placed under A-type conditions) illustrated in Figure S1. In doing so, we both relax the intense early-reproduction selection experienced by A-types and impose this same regime on C-types. Starting from these contrasting genetic backgrounds, we track phenotypic and genomic changes over several dozen generations to evaluate how populations respond to their new life-history cycles. If widespread balancing selection is maintaining functional variation in this system, we expect that: (1) evolutionary history will not impose strong, persistent constraints, and populations will converge genomically and phenotypically on long-standing counterparts, where under our design, this should translate into inverted genomic trajectories for the A→C and C→A given they are ultimately derived from the same ancestral population; (2) shifting A-types to C-type conditions will result in a widespread rebound in genetic variation, revealing low-frequency alleles that persisted despite the apparent loss of polymorphisms in past work, while the reciprocal C→A shift should produce a reduction in genetic variation; (3) replicate populations within each trajectory will show repeatable genomic responses, consistent with access to shared reservoirs of selectively maintained variation.

## Results

We analyzed two reciprocal selection trajectories, each replicated across 10 independently maintained populations. The long-established populations from which each trajectory was derived are referred to as the “founder populations”. We use “repeatability” to describe the consistency of evolutionary responses across replicate populations exposed to the same selection regime. We use “parallelism” to describe similar phenotypic or genetic changes arising in independently evolving populations, reflecting convergence on similar evolutionary outcomes.

### Adult Life-History Evolution: Mortality Trajectories

Founder populations exhibited the expected, long-established divergence in age-specific mortality between A-type and C-type selection regimes (Figs. S2,3; Tables S2-10), providing internal baselines for evaluating phenotypic change along reciprocal trajectories. At the time of the first mortality assay, C→A populations had experienced approximately 33 generations of early-reproduction selection, whereas A→C populations had experienced approximately 12 generations of delayed-reproduction selection. At this stage, both trajectory types differed significantly from their respective founders but exhibited asymmetric convergence toward the target selection regime consistent with differences in total number of generations under selection (Fig. S4A). C→A populations showed strong convergence with A-type founders across much of the adult lifespan, particularly at early adult ages, whereas A→C populations remained substantially divergent from C-type founders, showing early-phase mortality profiles (Tables S11-14).

At the next timepoint in the evolutionary trajectory, C→A populations had experienced approximately 143 generations of early-reproduction selection, whereas A→C populations had experienced approximately 51 generations of delayed-reproduction selection. Here we see both trajectory types have converged on the adult mortality phenotypes characteristic of long-term populations maintained under their respective selection regimes (Fig. S4B). Neither A→C nor C→A populations differed significantly from long-established populations of the same regime across adult life-history intervals, and both remained distinguishable from their original founders (Table S15-18).

To visualize systematic changes in adult mortality following reciprocal shifts in selection regime, we summarized mortality trajectories as the difference in mean instantaneous age-specific mortality rate between the evolved populations and their target-regime founder counterparts at each age interval, plotted as a function of age from egg. As selection proceeds, this difference is expected to approach but not necessarily reach zero, reflecting incomplete but ongoing evolutionary convergence between genetically distinct populations. In both reciprocal selection treatments, differences in age-specific mortality relative to the founder populations exhibit a smooth, wave-like pattern across the adult lifespan (Fig. 1). For A→C populations, there is convergence on mortality at early ages until a breakpoint day, at which point, mortality differences begin to increase gradually at successive ages. This breakpoint is pushed towards later ages with more generations of selection, indicating age-structured convergence toward C-type mortality profiles rather than uniform shifts across the lifespan.

**Figure 1.**
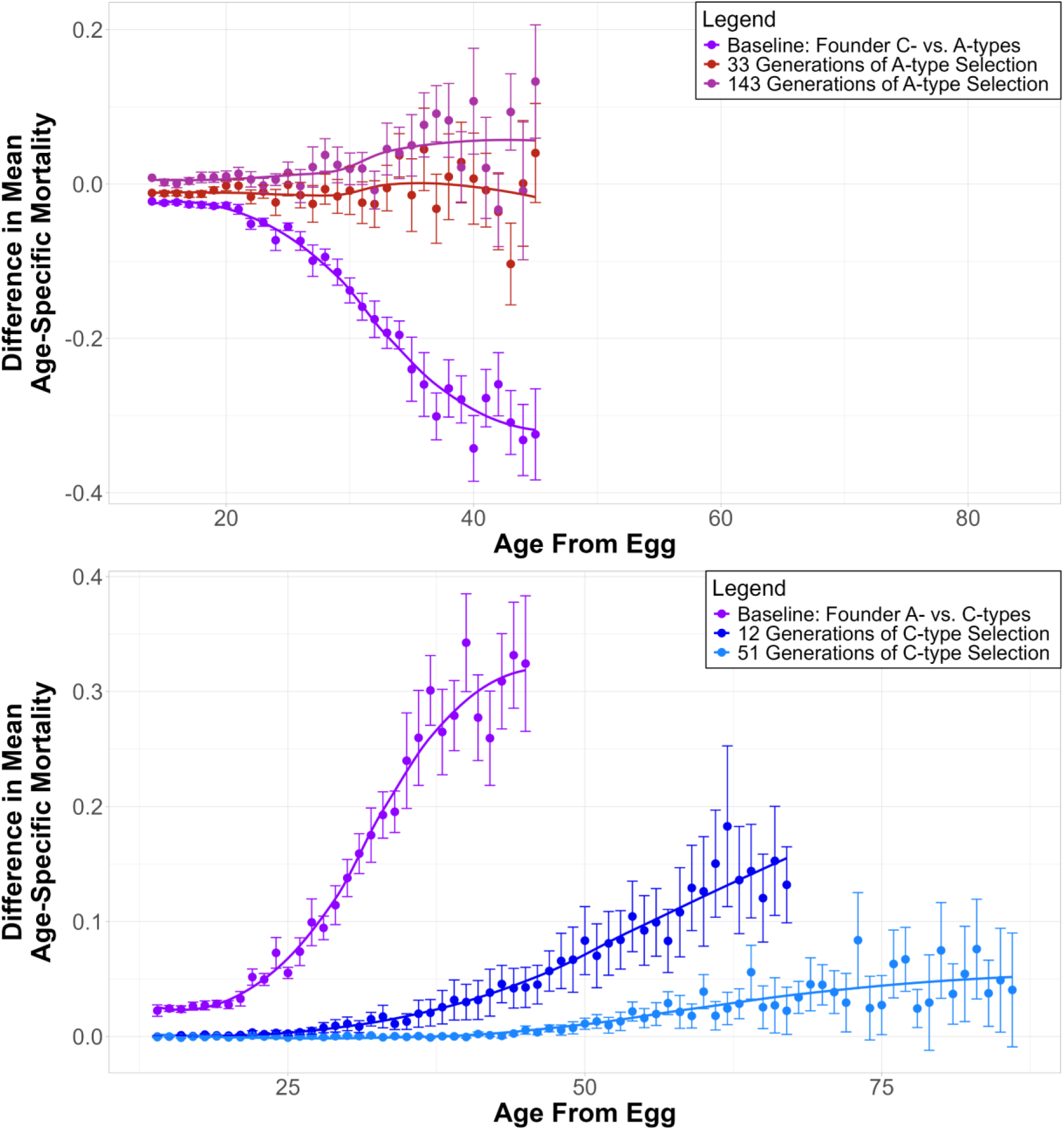
Waves of Convergence in selection. Differences in mean instantaneous age-specific mortality rate between population groups, plotted as a function of age from egg (days). Each point represents the mean difference across replicate populations at a given age interval; error bars indicate standard error across replicate populations. Lines show smoothed trends. The purple series in each panel shows the baseline mortality difference between the long-established A- and C-type founder populations, against which trajectory convergence is evaluated. (A) C→A trajectory: differences between C→A populations and A-type founder populations after ∼33 (magenta) and ∼143 (red) generations of early-reproduction selection. (B) A→C trajectory: differences between A→C populations and C-type founder populations after ∼12 (light blue) and ∼51 (dark blue) generations of delayed-reproduction selection.

### Developmental Timing Responses

To determine whether developmental timing responded concordantly to reciprocal life-history selection, we assayed pupation timing along the same evolutionary trajectories. For the first assay, A→C populations (∼6 generations of delayed-reproduction selection) and C→A populations (∼15 generations of early-reproduction selection) exhibited pupation timing intermediate relative to their respective founders (Figs. S5, S6; Table S19). At the next timepoint, A→C (∼64 generations), C→A (∼182 generations), pupation timing in both trajectory types converged on the phenotypes characteristic of long-term populations maintained under their current selection regimes (Fig. 2; Table S20). Thus, developmental timing shows a consistent and reversible response to reciprocal life-history selection, corroborating patterns observed for adult mortality.

**Figure 2.**
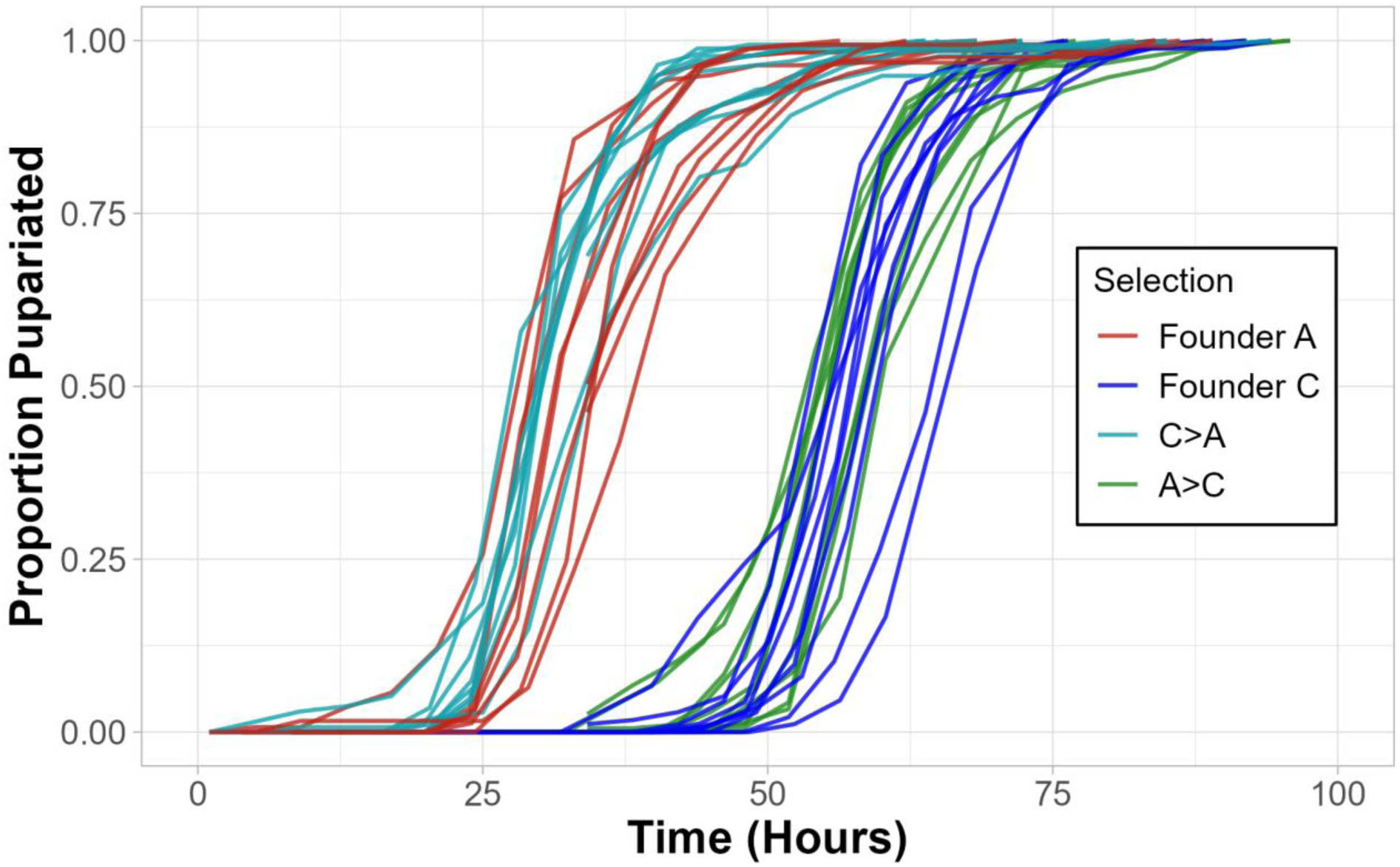
Cumulative pupariation timing, late phase. Cumulative proportion of individuals that have pupariated, plotted as a function of time from egg (hours) for founding populations A-type (red) and C-type (blue) and trajectory populations A→C (green) and C→A (teal). Each line represents an individual population replicate. At this late assay timepoint (∼64 generations of delayed-reproduction selection for A→C populations; ∼182 generations of early-reproduction selection for C→A populations), pupation timing in both trajectory types has converged on the phenotypes characteristic of long-term populations maintained under their respective selection regimes.

### Antiparallel Genomic Trajectories

To characterize genome-wide shifts associated with reciprocal selection, we performed a principal component analysis (PCA) on SNP frequencies from samples taken over the course of the experiment (Fig. 3). The first two components captured most of the variance, with PC1 explaining 48% and PC2 explaining 3.3% of genome-wide variation with no other individual PCs explaining more than 3%.

**Figure 3.**
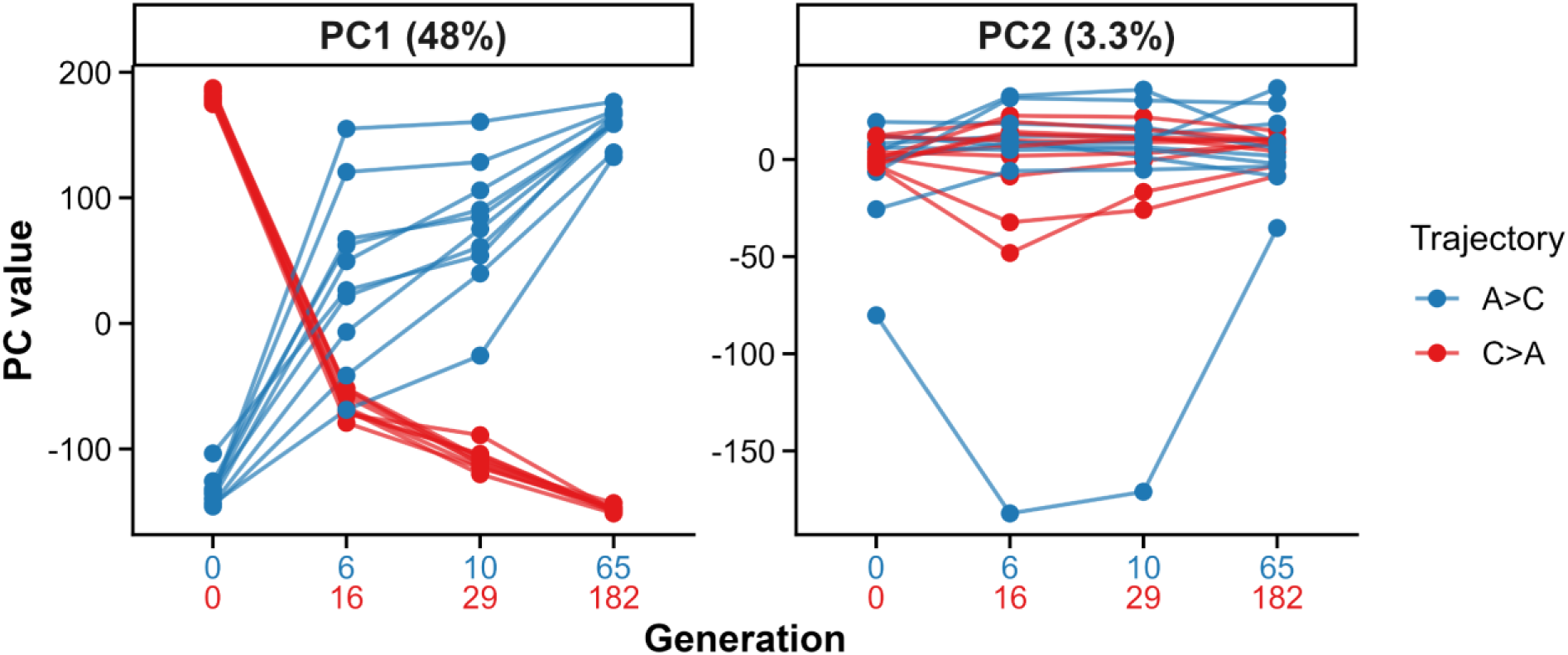
Antiparallel genomic trajectories under reciprocal selection. Lines indicate allele-frequency trajectories of replicate populations for A→C (blue) and C→A (red) across PC1 and PC2. Facet labels indicate the percentage of variance explained by each PC. Each line represents a replicate population, with points corresponding to successive generations (x-axis labels show generation numbers for each trajectory).

Projection onto PC1 revealed strongly antiparallel genomic trajectories. A→C populations shifted toward the cluster of long-established C-type populations, whereas C→A populations moved toward A-type populations along opposing directions. Dispersion among replicates was minimal at the final timepoints collected, whereas greater spread was observed among A→C populations at earlier stages relative to C→A populations, consistent with fewer generations having elapsed under delayed-reproduction selection.

To identify genomic axes that differ in their temporal response to selection, we analyzed the first ten principal components using linear mixed-effects models with selection treatment (A→C vs. C→A) and generation as fixed effects and population as a random effect. The treatment × generation interaction term tests whether PC scores change through time at different rates between treatments. A significant interaction was detected for PC1 only (Table S21: estimate = −4.19, Bonferroni-adjusted p = 2.13 × 10⁻⁶), indicating that PC1 scores change over generations at significantly different rates between the A→C and C→A trajectory groups, consistent with the opposing directional trajectories visible in Figure 3. No significant effects were detected for PC2 or any higher-order PCs.

To examine locus-specific responses, we quantified allele-frequency dynamics at all 1,389,963 individual SNPs using beta-binomial generalized linear mixed models with treatment, generation, and their interaction as fixed effects and population as a random effect. Under this framework, significant treatment × generation effects identify SNPs whose allele-frequency trajectory differs between the two regimes. We detected a widespread and highly polygenic genomic response to reciprocal selection (Fig. 4A), with 599,899 SNPs exhibiting significant treatment × generation interactions. Significance was determined based on two criteria: a permutation-based FDR threshold of 0.01 and a minimum effect size of 0.3 following an approach used by Kawecki et al. (2021). Conversely, no SNPs were significant with respect to the treatment or generation main effects based on these criteria. Among SNPs significant for the treatment × generation interaction term, the magnitude and direction of allele frequency change reveal that these responses are predominantly anti-parallel (Fig. 4B). Notably, the distribution is asymmetric: C→A populations exhibit larger allele-frequency shifts than A→C populations, reaching near-fixation at many loci, while A→C populations show more modest changes over their shorter experimental duration. Along with the PCA results, these findings demonstrate that reciprocal selection regimes produce extensive, genome-wide, and largely anti-parallel allele frequency changes.

**Figure 4.**
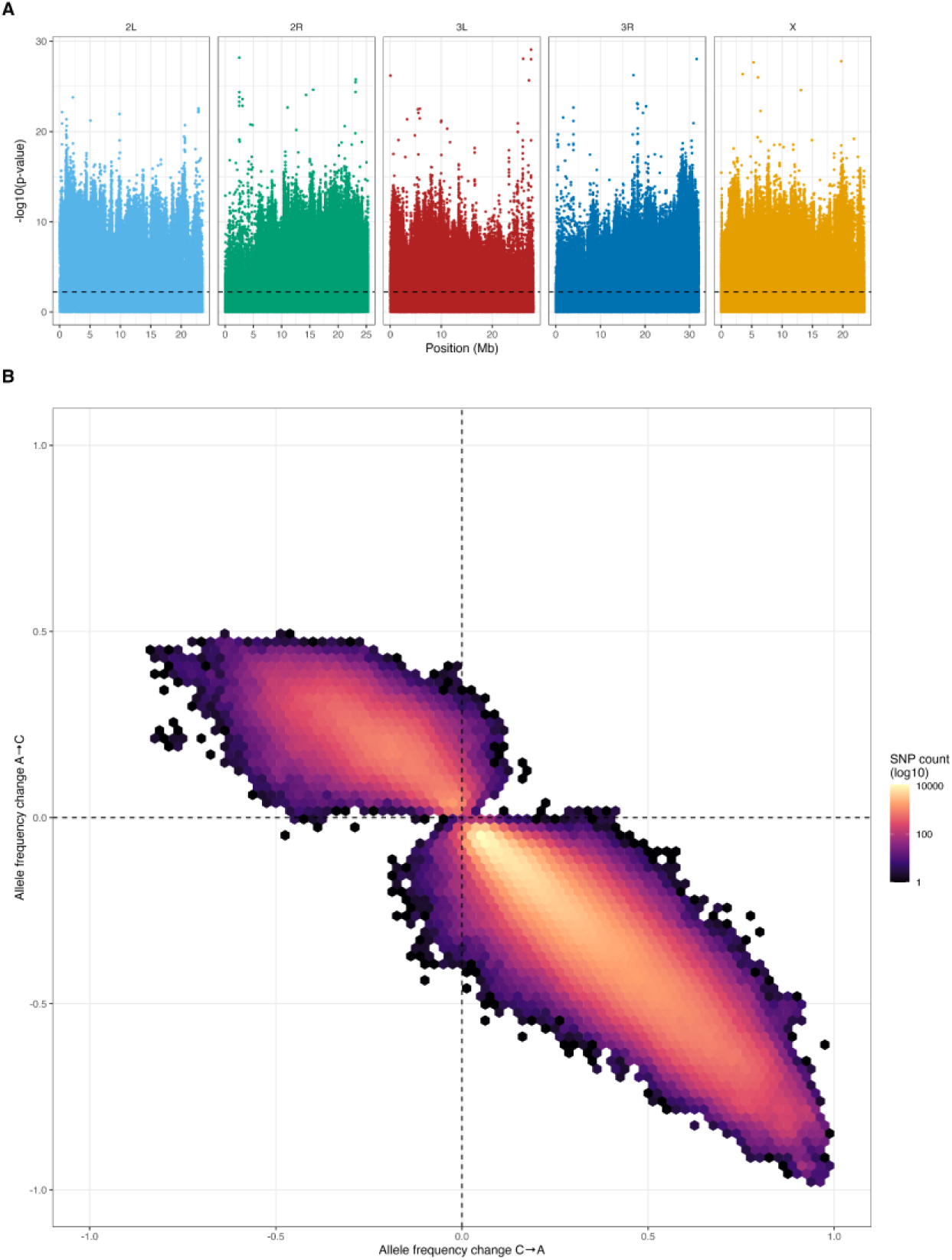
Genome-wide allele frequency responses to reciprocal selection. (A) Manhattan plot showing the significance of the treatment × generation interaction term from the beta-binomial GLMM across the five major chromosomal arms. The y-axis shows −log10(p-value) for each SNP. The dashed line indicates the permutation-based FDR and effect size threshold (FDR ≤ 0.01, frequency difference between trajectories ≥ 0.3). SNPs above the threshold show significantly different allele frequency trajectories between A→C and C→A populations over the course of the experiment. (B) Hexbin density plot showing the joint distribution of allele frequency changes between the first and last sampled timepoints for SNPs passing the FDR threshold, expressed relative to the A-allele. Positive values on the y-axis indicate allele frequency movement toward the A→C state; positive values on the x-axis indicate movement toward the C→A state. Density concentrated in the second and fourth quadrants indicate that the interaction term predominantly captures antiparallel allele frequency responses across reciprocal trajectories.

To highlight candidate loci for future investigation, we identified the most significant marker within each non-overlapping 10 kb window from the set of SNPs significant for the interaction term and annotated the 100 markers with the lowest p-values with respect to overlapping or proximal genes (Supplementary Table S24 and a full list of significant sites can be found on the Dryad directory associated with this project). Several biological themes emerged among the top candidates. A number of genes are involved in neural function and signaling (e.g., para, Ca-α1D, AstA-R2, Fmr1, Rim), consistent with the role of neural circuits in regulating reproductive timing and activity. A second group includes genes implicated in mitochondrial function and energy metabolism (e.g. TFAM, MICU1). We emphasize that this list is intended as a resource for hypothesis generation rather than evidence for any specific gene’s role. Given the highly polygenic nature of the response to selection in this experiment and confounded effects of linkage, our ability to make strong mechanistic claims is limited.

Lastly, to assess whether reversed populations had converged on the allele-frequency states of long-standing counterparts, we compared newly derived populations to the relevant founder populations (e.g. the final timepoint of the C→A trajectory versus extant A founders) that were sequenced at the end of the experiment using quasibinomial generalized linear models at each SNP (Fig. S7). Applying the same permutation-based FDR and effect size framework used in the trajectory analysis, we found no SNPs were significantly differentiated between newly derived and founder populations in either the A→C versus founder C comparison or the C→A versus founder A comparison. The absence of significant differentiation suggests that by the final timepoint, both trajectory types had converged on allele-frequency distributions statistically indistinguishable from those of long-established populations under the same regime.

### Relaxing Early-Life Selection Recovers Hidden Genetic Variation

Expected genome-wide heterozygosity, calculated on a per SNP basis then averaged across the genome, shifted in opposite directions along the two reciprocal trajectories (Fig. 5). A→C populations, which originate from low-heterozygosity A-type founders, showed a significant increase in heterozygosity over evolutionary time, whereas C→A populations, derived from high-heterozygosity C-type founders, exhibited a significant decline. These trends were consistent across replicate populations (n = 10 per trajectory) and were already evident at earlier generations. Comparisons between founder and endpoint populations confirmed highly significant changes in both trajectories (Welch two-sample t-tests with Holm-Bonferroni correction applied within trajectory, p < 10⁻¹¹). Together, these results demonstrate divergent genome-wide responses to reciprocal selection, with relaxation of early-life selection permitting recovery of genetic variation, while renewed early-life selection reduces heterozygosity.

**Figure 5.**
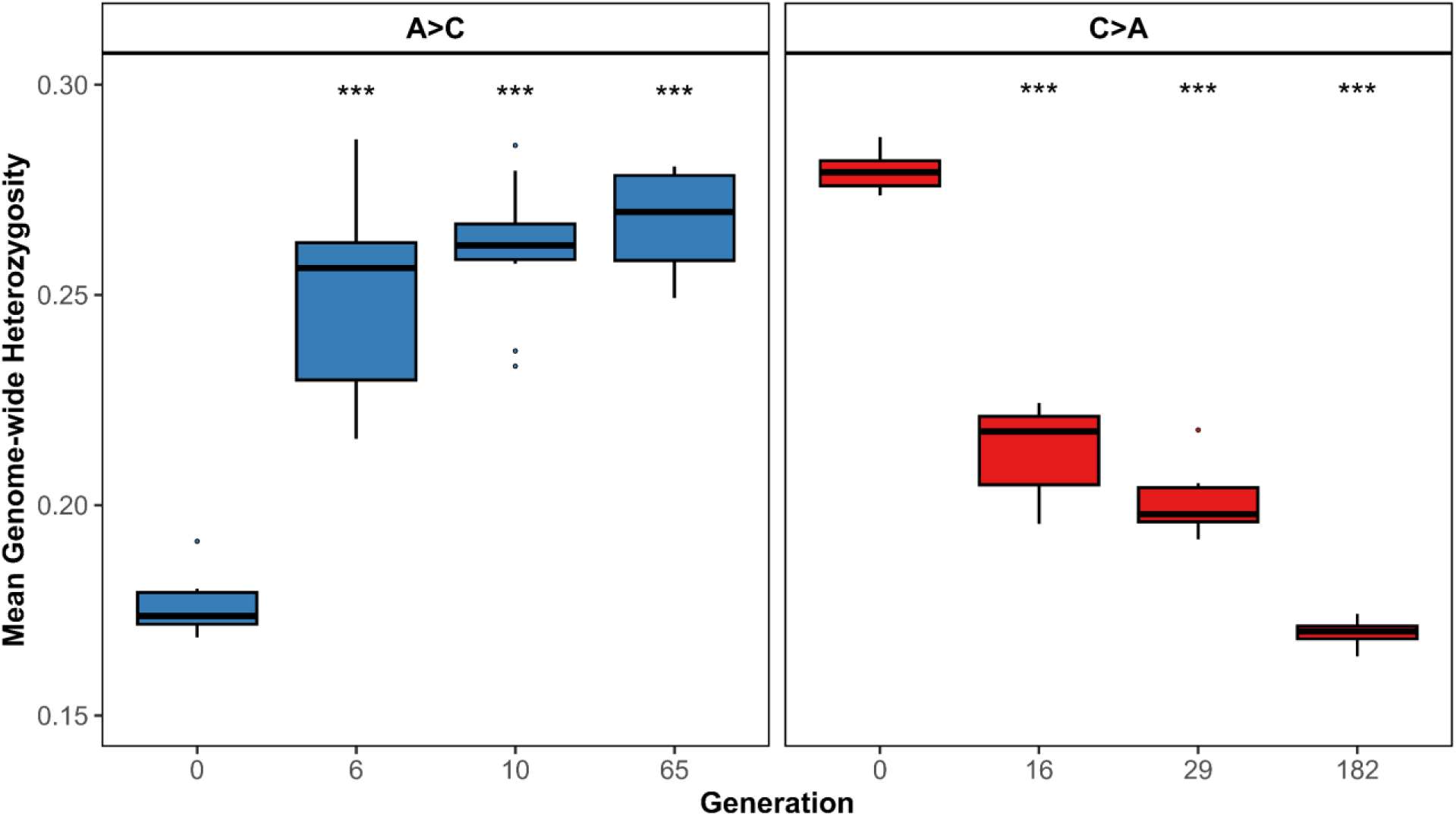
Genome-wide heterozygosity across generations for reciprocal selection trajectories. Mean genome-wide expected heterozygosity (H = 2f(1-f), averaged across all SNPs in the trajectory dataset) was calculated for each replicate population (n = 10 per trajectory) evolving under A→C (blue, generations 0, 6, 10, and 65) and C→A (red, generations 0, 16, 29, and 182) selection regimes. Boxplots show the distribution of mean heterozygosity across replicates at each sampled generation. A→C populations exhibit a rapid and progressive increase in heterozygosity following relaxation of early-life selection, whereas C→A populations show a consistent decline. Significance indicators reflect Welch two-sample t-tests comparing each post-founding generation to generation 0, with p-values adjusted using the Holm-Bonferroni sequential correction applied separately within each trajectory (* p < 0.05, ** p < 0.01, *** p < 0.001).

Many sites in the initial generation of the A→C trajectory appeared fixed under standard sequencing depth (∼90x), yet the genome-wide rebound observed following selection reversal suggested that this apparent fixation might instead reflect alleles present at frequencies too low to be reliably detected. To test whether these populations harbored hidden variation, we compared allele frequencies from a single founding population (A-type1, Timepoint 1) sequenced at higher depth (∼767x coverage) against sites appearing fixed in the standard-depth dataset. Specifically, we focused on sites that were identified as significant for the treatment × generation interaction term in the permutation-based FDR analysis (FDR < 0.01, effect size ≥ 0.3) and that showed either zero or full minor-allele counts under standard sequencing depth. Of the 599,899 SNPs meeting these criteria, 214,247 appeared pseudo-fixed in the A-type1 population. Deep sequencing revealed non-zero minor allele counts at 69,726 pseudo-fixed sites, while the remainder appeared fixed even at higher depth; subsequent analysis was performed on this subset

To minimize the influence of sequencing error, we applied a minimum minor allele count threshold of 3 prior to analysis. At a median coverage of ∼767x, a conservative Illumina per-base error rate of 0.001 predicts approximately 0.75 spurious reads per site, and sampling-with-replacement simulations based on a pool of 400 haploid alleles sequenced at ∼767x indicate a minimum detectable allele frequency of ∼0.00293; the count-3 threshold corresponds to ∼0.00391, providing a modest buffer above this detection floor. After filtering, 24,004 of the 69,726 pseudo-fixed sites (34.4%) retained detectable minor alleles. We examined the distribution of minor allele frequencies at these sites and compared it to a random subset of equal size drawn from the same deep-sequencing dataset (Fig. 6). While the random subset exhibited a broad range of allele frequencies (mean MAF = 0.175), pseudo-fixed sites were strongly enriched for low-frequency alleles, with a mean MAF of 0.016 and median MAF of 0.011. A permutation test (10,000 iterations) confirmed that the observed mean was significantly lower than expected under random sampling (Z = -158.59, p < 1 × 10⁻⁴), demonstrating that many sites appearing fixed under standard coverage in fact harbor rare alleles below standard detection thresholds.

**Figure 6.**
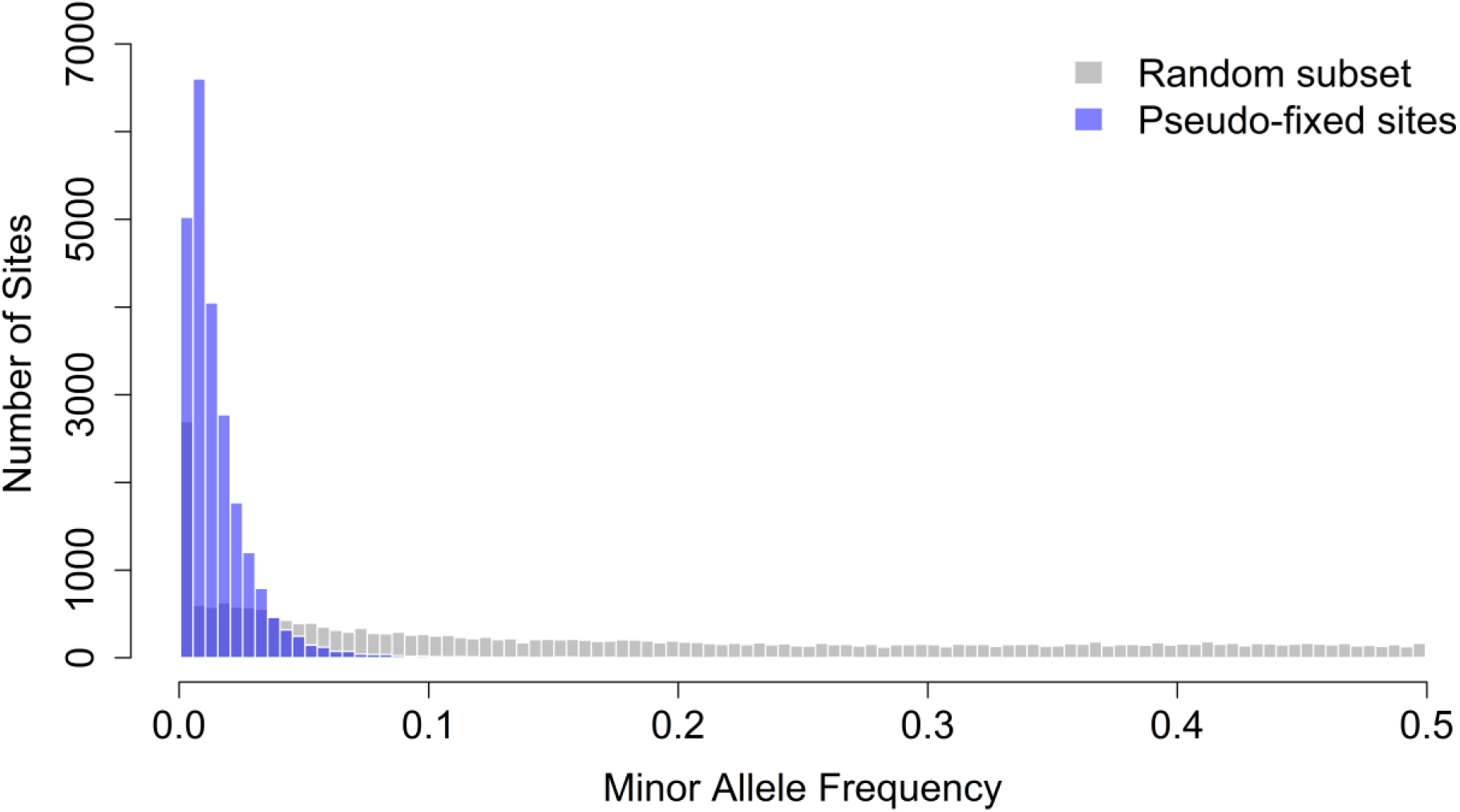
Deep sequencing reveals allele frequency variation at sites previously classified as fixed. Histograms depict the minor allele frequency (MAF) of SNPs in the A-type1 population sequenced at ∼767x coverage. Gray bars represent a random subset of SNPs equal in number to the pseudo-fixed sites identified in the trajectory dataset, while purple bars indicate sites classified as pseudo-fixed under standard coverage (permutation-based FDR < 0.01, effect size ≥ 0.3, treatment × generation interaction term). Pseudo-fixed sites are overwhelmingly skewed toward low-frequency alleles, indicating that these loci appear fixed under standard coverage but retain hidden variation at higher sequencing depth. Permutation testing (10,000 replicates) confirmed that the observed mean MAF of pseudo-fixed sites (0.016) is dramatically lower than expected under random sampling (mean null = 0.175, SD = 0.001; Z = -158.59, p < 10⁻⁴), demonstrating that these pseudo-fixed sites represent a significant depletion of minor alleles relative to genome-wide expectations.

To assess whether this low-frequency reservoir could be explained by neutral drift alone, we estimated the effective population size (Ne) of each A-type founder population from empirical allele frequency changes observed over approximately 219 generations, using the temporal method of Jorde and Ryman (1995) as implemented in the poolSeq package (Taus et al. 2017). Ne estimates across the 10 replicate populations ranged from approximately 338 to 1,270, with a mean of approximately 897 and a median of approximately 987, consistent with the census size of ∼1,500 individuals maintained per population (Table S22). At these effective sizes, alleles present at frequencies on the order of 0.001 face a high per-generation probability of loss to drift, confirming that the low-frequency reservoir detected by deep sequencing is unlikely to persist passively over the ∼1,000-generation history of the A-type populations. The consistency of this reservoir across 10 independent replicate populations subject to equivalent drift dynamics argues instead for active maintenance through balancing selection.

Of the 599,899 SNPs with significant treatment × generation interactions, 38,227 (6.37%) had zero minor allele counts across all 10 A-type founder populations under standard sequencing depth, identifying them as members of the pseudo-fixed hidden reservoir. The remaining 93.6% were already segregating at detectable frequencies in the A-type founders, with a mean founder MAF of 0.203, indicating that most of the genomic response to selection reversal drew on variation already present at appreciable frequencies. The emergence of hidden alleles across replicate populations was strikingly consistent: all 38,227 pseudo-fixed responders became detectable in at least 6 of 10 A→C populations by generation 65, 99.2% emerged in 8 or more replicates, and 71.6% emerged in all 10 (mean = 9.65 replicates). This degree of parallelism across independently evolving populations is inconsistent with stochastic emergence and instead confirms that these alleles were present at cryptic frequencies in the founders and responded reproducibly to the reversal of selection (Table S23).

By generation 65, pseudo-fixed responders reached a mean frequency of 0.254 and median frequency of 0.225 across replicate populations, with even the least responsive sites (10th percentile) climbing to 0.152. This rapid rise from cryptic frequencies below standard detection thresholds to population-level frequencies exceeding 0.20 within 65 generations of selection reversal underscores both the strength of selection acting on these alleles and the biological reality of the hidden reservoir detected by deep sequencing. The 100 highest-ranked lead SNPs from the interaction term analysis, selected by FDR and then by p-value from the set of most significant markers per 10 kb window, along with their allele frequency trajectories and cross-replicate consistency, are provided in Supplementary Table S24.

## Discussion

Long term experimental evolution studies have repeatedly shown that adaptation in sexually reproducing populations is dominated by modest, coordinated allele frequency shifts across many loci rather than fixation at single large-effect variants (Burke 2012; Barghi et al. 2019; Barghi and Schlötterer 2020). While this “shifts not sweeps” pattern is now well established, deeper questions emerge regarding the mechanisms responsible for maintaining such extensive variation, especially in cases with sustained and intense selection. In the absence of frequent de novo mutations or environmental fluctuations, classical theories predict that drift and directional selection should steadily erode standing genetic variation as populations converge on a new optimum. In this study we demonstrate that genetic variation persists and remains evolutionarily accessible even after hundreds of generations of intense life-history selection, even from populations associated with major loss of variation (Graves et al. 2017). These results provide strong evidence for the active maintenance of polymorphisms through balancing selection.

### Reciprocal selection reveals active maintenance of genetic variation

A central result of this study is the rapid phenotypic reversibility of populations with opposing selection regimes. Adult mortality and developmental timing assays demonstrated clear responses to selection, with populations evolving toward the characteristic phenotypes of long-established A- and C-type populations. Crucially, this reversibility was observed even in A-type populations, which are characterized by substantial genome-wide reductions in genetic variation. These results suggest that alleles contributing to delayed reproduction, extended lifespan, and slower development were actively maintained at low frequencies in the A-type populations in some manner.

Importantly, phenotypic reversibility was mirrored at the genomic level. Principal component analysis reveals a global anti-parallel response with A→C and C→A populations moving in opposing directions toward their respective selection state. When we examined locus-specific responses using beta-binomial generalized linear mixed models, the interaction term accounted for all significant SNPs, identifying loci whose allele frequency trajectories differed between regimes over time. As shown in Figure 4B, the large majority of these represent antiparallel responses, confirming that global antiparallel trajectories were not diffuse or idiosyncratic. Rather, they appear to be driven by repeatable and parallel shifts in target sites and linked regions. In total, these results are consistent with widespread balancing selection ensuring populations consistently had access to the same subsets of standing variation when selection was reversed. These results build on those of Graves et al. (2017), showing that in this system recent selection regime is the primary determinant of a population’s genetic state, even when prior selection regimes involved hundreds of generations of very intense selection.

### Relaxation of early-life selection reveals hidden variation

Because A-type populations are characterized by substantial genome-wide depletion of genetic variation, the wholesale rebound in heterozygosity observed in the A→C populations suggest these apparent losses are not as complete as they otherwise seem. Deep sequencing of an A-type population reveals that many seemingly “fixed” sites instead harbor minor alleles below standard detection thresholds, indicating that when selection is relaxed, these alleles are able to respond and produce a genome-wide rebound in heterozygosity. Although this inference is based on deep sequencing from a single replicate population, the consistent rebound observed across independent lines argues that low-frequency reservoirs are a general feature of A-type populations and help explain their rapid genomic and phenotypic recovery. Whether this persistent low-frequency variation is introduced or actively maintained warrants careful consideration.

Potential sources of newly introduced variation are limited. Mutation alone is an insufficient explanation, as census sizes are too small to produce coordinated allele-frequency shifts across thousands of loci within only a few generations. Accidental migration between independently maintained stocks is equally implausible given the required rate of exchange. Prior simulation work in Phillips et al. (2016) demonstrates that unrealistically high and persistent migration rates would be required to produce the levels of similarity observed between replicate populations in this system. Such rates are inconsistent with independently maintained laboratory stocks where migration is accidental by definition. The reproductive windows of the regimes further preclude systematic gene flow between groups: C-type flies are still in the pupal stage during the A-type flies reproductive window, and at the ages when C-types reproduce, A-types are competitively inferior. Moreover, for migration to generate the highly parallel rebound we observe, identical alleles would have to enter multiple replicate populations at roughly the same time, persist at low frequencies despite being disfavored, and then rise in concert when selection shifts.

Having excluded plausible sources of newly introduced variation, the remaining explanation is that this reservoir is actively maintained. Importantly, the deep sequencing data provide a conservative view: they capture only alleles that have survived to the point of sampling, whereas any variants lost to drift during the ∼1,000-generation history of the A-type populations are necessarily invisible. The observed reservoir is therefore not a transient accumulation of rare alleles, but a filtered subset that has repeatedly persisted through the combined effects of drift and strong directional selection. Under the estimated effective population sizes for this system, such persistence is not expected under neutrality. Drift should steadily erode variation across the low-frequency spectrum, including alleles at the median detected frequency of 0.011, making their long-term survival across ∼1,000 generations unlikely in the absence of opposing forces. Under a purely deleterious model, alleles disfavored in A-type populations would be expected to reach mutation-selection balance frequencies far below what standard sequencing could detect, and their distribution across replicate populations would reflect stochastic mutation input rather than a shared reservoir. The observed frequencies and cross-replicate consistency are inconsistent with this expectation and instead point to alleles held above mutation-selection balance by opposing selective forces.

The pattern of response following selection reversal further strengthens this inference. The same alleles emerged reproducibly across independent replicate populations, with the vast majority becoming detectable in 8 or more replicates and over 70% appearing in all 10. Moreover, the magnitude of this response is striking: pseudo-fixed alleles rose from cryptic frequencies below standard detection thresholds to a mean of 0.254 across replicate populations within just 65 generations, a trajectory inconsistent with drift at the Ne values estimated for this system and indicative of strong positive selection following regime reversal. This pattern of coordinated and rapid re-emergence is difficult to reconcile with stochastic processes alone. Instead, it implies that these alleles are maintained by consistent selective pressures acting across populations, most parsimoniously in the form of widespread balancing selection that preserves a reservoir of low-frequency variation available for rapid and repeatable response when the selective regime changes.

### Highly repeatable and parallel genomic responses argue against historical contingency

A striking feature of the reciprocal trajectories is the tight clustering of replicate populations at our final collection points. Despite originating from similar yet distinct founders and evolving along opposite trajectories, populations subjected to the same selection regime converge on highly reproducible genomic states within their respective regimes. The limited number of loci that remain significantly differentiated between reciprocally selected populations and their corresponding founders further underscores the repeatability of these evolutionary outcomes. Importantly, the small asymmetry observed between the A→C and C→A comparisons is readily explained by differences in the number of generations experienced under selection, rather than by intrinsic limits on reversibility. These results argue strongly against a dominant role for historical contingency or idiosyncratic paths in shaping long-term genomic outcomes. Instead, they indicate that selection repeatedly draws on the same reservoirs of standing variation and drives populations toward predictable genomic outcomes.

Next, the largely parallel genomic responses within each reciprocal trajectory clarify how selection acts on standing variation in this system. Even under stringent significance thresholds, a substantial fraction of the genome shifts in both A→C and C→A trajectories, highlighting the pervasive impact of altered life-history selection.

### Current trajectories support robust inference despite ongoing adaptation

Collectively, our data suggest that the A→C populations are perhaps still adapting, and our conclusions therefore reflect the current direction and consistency of the response rather than a completed evolutionary endpoint. Nevertheless, the concordance of phenotypic reversibility, genome-wide antiparallel allele-frequency shifts, rebounds in heterozygosity, and agreement across multiple genomic analyses indicates that the central features of the response are already well established. While additional timepoints may refine the magnitude of these changes, they are unlikely to qualitatively alter the interpretation supported by the patterns already observed.

### Antagonistic pleiotropy as a plausible mechanism

Taken together, these results are consistent with the antagonistic pleiotropy models developed for discrete-generation populations in Rose (1982). Importantly, our interpretations do not require that individual fitness-components exhibit classical overdominance or frequency-dependent selection. Rather, antagonistic pleiotropy distributed across many fitness-components can collectively generate balancing dynamics at the level of net fitness. In such a system, fixation is constrained because alleles that are favored under one fitness component impose costs on others, preventing any single genotype from achieving universally superior fitness across the full age-structured reproductive schedule. This population-level resolution of trade-offs naturally leads to the maintenance of extensive genetic variation, even under constant environmental conditions. Because the Rose laboratory’s A and C regimes impose selection directly on reproductive timing, they inherently act on age-structured fitness, a core context in which antagonistic pleiotropy is predicted to generate life-history trade-offs (Rose 1985).

It is worth noting, however, that antagonistic pleiotropy may primarily operate by reducing net selection coefficients rather than generating the kind of stable frequency-dependent equilibria typically associated with canonical balancing selection. If so, individual allele fates may still be governed substantially by drift, particularly at the Ne values we estimate for this system. The classic theoretical consensus, formalized by Hedrick (1999), holds that stable polymorphism under antagonistic pleiotropy is largely confined to cases where selection coefficients are large and roughly symmetric, making it a theoretically restricted mechanism. However, this notion has recently been challenged by Brud and Guerrero (2026), who conducted a geometric analysis of the full parameter space across six models of antagonistic selection and found that the region permitting protected polymorphism is substantially broader than previously appreciated when the assumption of constant dominance across fitness components is relaxed. Non-reversing dominance alone contributes at least 25% of the stabilizing parameter space across all models, and the potential for polymorphism remains substantial even under fairly weak selection and asymmetric selection coefficients. Density-dependent selection with respect to genotype represents an additional independently plausible mechanism that could generate negative frequency dependence and maintain variation, and we cannot exclude its contribution in this system. Antagonistic pleiotropy therefore remains a working hypothesis, consistent with the broader theoretical landscape but not yet established as the operative mechanism.

Empirical support for the biological plausibility of this hypothesis also comes from Zwoinska et al. (2026), who conducted an evolve-and-resequence experiment in Callosobruchus maculatus and found that relaxing sexually antagonistic trade-offs released large, repeatable allele frequency shifts at loci that also carried signatures of long-term balancing selection in the ancestral population, while intensifying antagonism produced genomic responses indistinguishable from a relaxed selection control. This dynamic closely parallels what we observe in our system, where relaxing early-life selection in A-type populations releases a reservoir of low-frequency variation in a repeatable and parallel manner across independent replicates. Taken together, the theoretical breadth identified by Brud and Guerrero (2026) and the empirical parallels from Zwoinska et al. (2026) suggest that antagonistic pleiotropy, while not the only plausible mechanism, represents a well-grounded and empirically motivated explanation for the patterns we report.

## Conclusion

Broadly, these findings contribute to a growing body of evidence that adaptive evolution in sexually reproducing populations frequently proceeds through coordinated, genome-wide shifts in allele frequencies rather than classic hard sweeps (Burke 2012; Barghi et al. 2019; Barghi and Schlötterer 2020). In our reciprocal selection framework, the largely antiparallel responses across replicate populations and chromosome arms, together with the capacity for directional reversal under altered age-specific selection, indicate that substantial genetic variation is maintained over long evolutionary timescales. Rather than being exhausted by prolonged selection, this standing variation remains available to fuel rapid and repeatable responses when the selective regime changes. Such patterns serve as strong signatures of balancing processes operating across many loci, maintaining polymorphism through time and enabling predictable yet reversible genomic trajectories. Viewed in this light, our results contribute to the growing body of literature on the importance of considering balancing selection in evolutionary genetics, not as a marginal phenomenon, but as a major driver of observed patterns of genetic variation (Kawecki et al. 2021, Zwoinska et al. 2026). More specifically, these findings have potential implications for conservation genomics, where loss of genetic diversity is often interpreted as evidence of extinction debt (Gargiulo et al. 2025). In systems where selection actively maintains variation, alleles appearing lost under standard sequencing may persist at cryptic frequencies and respond when selective conditions change, warranting caution in assuming that apparent diversity loss is irreversible in all contexts.

## Materials and Methods

### Experimental Populations

All *Drosophila melanogaster* populations used in this study trace to the IV population, mass-sampled in 1975 from the wild South Amherst population described by Ives (1970) and maintained as a large outbred laboratory strain under standard banana agar media. The B and O selection treatments were derived from IV in 1980 (Rose 1984; Rose et al. 1992), and from these, multiple experimental evolution lines have been derived, with five independently replicated populations subjected to distinct selection regimes. Of these treatments, the laboratory cultured ten early-reproducing A-types and ten late-reproducing C-types, achieving hundreds of generations of sustained selection (Chippindale et al. 1997), demonstrating strong convergence across life-history characteristics and genomics (Burke et al. 2016; Graves et al. 2017). Given this strong convergence within each regime, we refer to these two sets of 10 populations as the founder A-types and the founder C-types respectively.

Building upon this work, we derived 20 new populations from the progenitor A-types and C-types. The founder populations themselves were not subjected to any change in selection regime and have been maintained continuously under their original A-type and C-type conditions throughout the duration of this experiment, serving as stable genomic and phenotypic references. The new populations were created by imposing the reverse selection on offspring collected from the founders: A-type selection was imposed on 10 populations derived from the progenitor C-types to create the C→A-types, and C-type selection was imposed on 10 populations derived from the progenitor A-types to create the A→C-types.

C→A-type populations were created by collecting eggs from the 10 founder C-types, which continued unaltered on their 28-day C-type regime, and only permitting the first ∼20% developing flies to contribute to the next generation. This was accomplished by rearing eggs from each generation in carousels, which feature food caps housed in a dome to contain the earliest developing flies within until enough individual flies have eclosed to create a new outbred population. The earliest derived flies contained in the carousel were then gassed with CO2 and pooled into Plexiglas cages to lay eggs for the next generation. Multiple carousels were utilized per population to yield large cohort sizes to prevent inbreeding depression from the intense selection regime. With each vial containing 80+ eggs, 80 vials in each carousel, and ∼3 carousels per population, ∼19,200 total eggs were obtained each early generation, with approximately ∼3,840 young adults constituting the top 20% of early developers permitted to create the next generation. With each successive generation of directional selection, average development time decreased, reducing the need for additional carousels and thereby minimizing the risk of large population bottlenecks as the experiment continued. Once the C→A-type populations were reliably reproducing on Day 10 without complications, they were maintained like their progenitor A-type counterparts with only vials and cages, and no carousels.

A→C-type populations were created by collecting eggs from the 10 progenitor A-type populations, which continued unaltered on their 10-day A-type regime, and allowing only those females still alive and fecund by day 26 to contribute to the next generation. Because A-type populations have substantially shorter adult lifespans than C-type populations, additional cohort cages were required to ensure that a sufficient number of flies survived to day 26 to propagate the next generation. This involved collecting ∼7,500 eggs for each population, transferred on day 14 into 5 separate cages containing ∼1,500 individuals. Standard banana media was fed to the cohorts every other day until day 26, when cages were condensed to ensure enough living and breeding individuals were pooled to maintain at least ∼1,500 individuals to produce the next generation for a given population. Intense forward selection continued with multiple cohorts collected from separate cages and pooled within a population cage to elevate census sizes until populations could reliably be maintained with a single cage.

### Data Collection

At several intervals, phenotypic assays and genomic samples were collected to track the two opposing trajectories as their constituent populations adapted to their new selection regimes. By measuring these newly derived populations at multi-generation intervals, we were able to map the initial trajectories of phenotypic and genotypic change in response to selection. Although phenotypic assays and genomic sampling were conducted at the same chronological time points, A→C and C→A trajectories accumulated different numbers of generations due to differences in generation time and are therefore treated as distinct experiments.

### Adult Mortality

Adult mortality assays followed the protocols described in Burke et al. (2016), with minor modifications. Progenitor stock populations were sampled on a 14-day culture cycle for two generations, staggered by one day to align treatments on a shared calendar and minimize environmental variation. Each population was represented by two cohorts (e.g., A-type1α and A-type1β) of approximately 1,500 flies housed in Plexiglas cages and supplied daily with fresh banana–molasses medium (Phillips et al. 2018) supplemented with yeast to induce oviposition after transfer on day 14 from egg. At the same time each day, all dead flies were removed, sexed, and recorded for each cohort until no individuals remained. Mortality data were pooled across the two cohorts to obtain daily death counts for each population.

The Burke et al. (2016) protocol was modified to eliminate the use of carbon dioxide for condensing cohorts when census densities were low. Instead, flies remained in their original cages for the duration of the assay, with internal cleaning performed as needed to remove waste accumulation from the cage walls. Adult mortality assays were conducted on founder populations and on the selected populations at multiple points along the adaptive trajectory, corresponding to approximately 12 and 51 generations for A→C populations and approximately 33 and 143 generations for C→A populations.

Age-specific mortality rates were analyzed using linear mixed-effects models fit independently at each 3-day age interval. For significance testing, log-transformed instantaneous mortality rate was modeled as a function of selection regime as a fixed effect, with population nested within age as a random effect. Age intervals were included only when all expected replicate populations contributed observations, ensuring balanced comparisons. Models were fit using the lme function from the nlme package in R, with pairwise comparisons conducted between selection regimes. Multiple comparisons within each pairwise test were corrected using a Bonferroni threshold applied across the number of age intervals tested. All analyses were conducted on female mortality only. To visualize the wave of convergence pattern, mean differences in age-specific mortality between trajectory and founder populations were additionally calculated on the untransformed scale and plotted as a function of age. Results from all pairwise comparisons are summarized in Tables S2--S18.

### Pupariation Timing Analysis

Each larval development assay was conducted following two generations of standardized rearing. Ancestral founder populations (A-type and C-type) and trajectory populations from the C→A and A→C treatments were assayed for pupation timing. For each population, eggs were collected on nutrient-free agar plates, and 60 first-instar larvae were individually transferred to food vials, with three vials established per population. Vials were maintained in a sealed incubator at 25 °C under constant light conditions. Larval development was monitored every four hours for 14 days, and the time of pupariation was recorded for each individual upon detection of newly formed pupal casings. To minimize temporal bias and blocking effects among treatments, one replicate population was initiated per day; however, once established, all replicates were monitored at identical time intervals throughout the assay.

Pupation data were analyzed using linear mixed-effects models to evaluate the effects of selection regime on developmental timing. For each vial, the proportion of individuals pupated was modeled as a function of selection regime, time, and their interaction, with vial nested within replicate as a random effect to account for non-independence among larvae sharing the same vial. Replicate identity corresponded to the day of assay initiation and was included to account for blocking effects. For each vial we have assayed the proportion of developed flies *f*_*hijk*_, from selection regime *(h* = 1,2), time in hours (*i* = 4,8, . . . *T*), replicate (*j* = 1,2, . . .5), and vial (*k* = 1,2,3). Models were fit separately for each pairwise comparison among selection regimes, with *h* = 1 serving as the reference group and *h* = 2 as the comparison group; the specific populations assigned to each role varied by comparison. For the early-phase assay, three pairwise models were fit per dataset: founder A-type vs. founder C-type, founder C-type vs. C→A, and C→A vs. founder A-type. An analogous set of three comparisons was fit for the A→C dataset substituting A→C populations in place of C→A. For the late-phase assay, five pairwise models were fit: founder A-type vs. founder C-type, C→A vs. A→C, all A-type vs. all C-type, C→A vs. founder A-type, and A→C vs. founder C-type. The model can be summarized as:

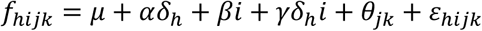

where *δ*_*h*_ = 0 if *h* = 1 and 1 otherwise.

- *⍺* is the fixed effect of selection,
- *β* is the fixed effect of time,
- *γ* is the fixed effect for the interaction of time and selection regime
- *θ*_*jk*_ is the random effect of vial *k* nested within replicate *j*
- *ɛ*_*hijk*_ is the residual error

In short, Proportion Developed = μ + Selection Regime + Time + Selection Regime × Time + Replicate/Vial + ε. Replicate refers to reps 1, 2, 3, 4, or 5 and corresponds to day of experiment start to account for blocking effects. Pupariation timing assays were conducted at two points along the adaptive trajectory. The early-phase assay corresponded to approximately 15 generations of selection for C→A populations and approximately 6 generations for A→C populations. The late-phase assay corresponded to approximately 182 and 64 generations of selection for C→A and A→C populations, respectively.

### Genomic Sampling

Samples for genomic analysis were collected following the protocols described in Graves et al. (2017). For each population, approximately 1,500 adults per population were maintained in separate Plexiglas cohort-cages with banana–molasses medium. Prior to sampling, populations underwent two lead-in generations on a standardized 14-day culture cycle, with collections staggered by one day among cages to minimize environmentally induced variation that could bias allele frequency estimates. This 14-day common garden interval was chosen to standardize sampling age between the regimes: it precedes the accelerated mortality of A-type adults while ensuring that C-type adults have fully eclosed, avoiding bias from incomplete representation of the cohort. Flies were anesthetized with CO₂ before sampling, and each sample consisted of a pool of ∼200 randomly collected females per population. Each replicate population was sampled and sequenced independently as a separate pool, preserving the replicated structure of the experimental design and enabling population as a unit of replication in all downstream analyses. The ten A-type and ten C-type founder populations were sequenced at time points 1 and 4, and the ten A→C and ten C→A reciprocal populations were sampled at time points 2, 3, and 4.

### DNA Extraction & Sequencing

Pooled samples of ∼200 female flies were immediately flash-frozen in liquid nitrogen and stored at −80°C prior to genomic extraction. Samples were homogenized in liquid nitrogen prior to extraction using Qiagen Puregene kits, and DNA concentration was assessed using a NanoDrop spectrophotometer. Extracted DNA pools were submitted to the UC Irvine Genomics Research and Technology Hub (GRTH) for gDNA library construction and sequencing. Samples were mechanically sheared to a target size of ∼150 bp, and an AMPure XP bead cleanup (1.8x) was performed on a subset of samples exhibiting smaller fragment profiles indicative of partial degradation; all samples were verified by Bioanalyzer prior to library construction. Each replicate population received a unique dual-index (UDI) barcode (IDT UDI indices), and libraries were sequenced on an Illumina NovaSeq X Plus using 150 bp paired-end chemistry (PE150). To avoid index collision, samples from different collection years were multiplexed on separate lanes. Samples were initially sequenced at a target depth of ∼30 million read pairs (∼50x average genome coverage), then re-sequenced to bring the combined total to a target depth of ∼60 million read pairs (∼100x average genome coverage). One population (A-type1, Timepoint 1) was additionally subjected to deep sequencing at a target depth of 500 million paired reads; library concentration was re-verified by Qubit prior to this run.

#### SNP Data Processing

Raw sequencing data were received from the sequencing center as ∼150 bp paired-end FASTQ files. All 100 standard sequencing samples were sequenced across multiple runs to a combined target depth of ∼60 million read pairs (∼100x coverage). One population (A-type1, Timepoint 1) was additionally subjected to deep sequencing at a target depth of 500 million read pairs on a dedicated lane; library concentration was re-verified by Qubit prior to this run, and this sample was kept on a separate lane from standard samples to avoid index collision. Reads were mapped to the *Drosophila melanogaster* reference genome (dm6, including the ISO1 mitochondrial genome) using Novoalign v4.04.01 (Novocraft Technologies 2025). Alignment was performed in paired-end mode (-r RANDOM, -I PE 250, 50) with default settings. A homopolymer filter was used to exclude low-complexity reads and reads failing to meet quality thresholds were removed prior to alignment.

Across standard sequencing samples, mean unique alignment rate was 82.5% (range 71.7--86.1%), with a mean proper pair rate of 81.4%. Multi-mapped and unmapped reads comprised 8.9% and 7.6% of total reads respectively, and approximately 1% of reads were removed by the homopolymer filter. For the deep sequencing sample, unique alignment rate was 81.6% and proper pair rate was 86.3%. The “Novosort” function was then used to sort the resulting BAM files and identify PCR duplicates. BAM files corresponding to separate sequencing runs for a given population were merged using Bamtools (Barnett et al. 2011). Merged BAM files for individual samples were then combined in mpileup format using SAMtools (Li and Durbin 2009). PoPoolation2 (Kofler et al. 2011) was then used to convert this mpileup to a “sync” file for ease of processing. Lastly, a Python script was used to convert the sync file into a SNP table with minor allele counts and total coverage for all polymorphic sites for each population in the dataset.

Only biallelic sites were considered, and the minor allele was defined as the less common nucleotide across all samples. We filtered with a minimum 20x and maximum 500x coverage in each sample, and polymorphic sites were defined as those with a minimum minor allele frequency of 0.02 across all samples. This resulted in a dataset of 1,389,963 SNPs. For standard sequencing samples, mean SNP coverage per sample was 90.2x (median 90.7x), ranging from 63.1x to 114.8x across samples. Coverage was consistent across population types and collection timepoints, indicating that differential coverage is unlikely to confound allele frequency comparisons between A→C, C→A, and founder populations. For the deep sequencing sample, mean SNP coverage was 767x (median 762x), with the central 90% of sites ranging from 483x to 983x across sites. Full sequencing and coverage statistics for both standard and deep sequencing are provided in Table S25.

#### Principal Component Analysis

To summarize genome-wide allele frequency variation, principal component analysis (PCA) was conducted on SNP frequency matrices using the prcomp function in R (R Core Team 2024). Data were mean centered but not scaled, as allele frequencies are already on a common scale and scaling to unit variance would inappropriately upweight low-variance sites. To examine the evolutionary trajectories of populations over time, each PC (PC1–PC10) was modeled as a function of trajectory and generation using linear mixed-effects models:

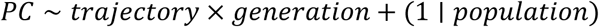

where *trajectory* represents the evolutionary direction (A→C or C→A), generation is a continuous variable representing generations since the start of selection, and population is a random intercept accounting for replicate-specific effects. Models were fit using *lmer* from the lme4 package, and significance of fixed effects was assessed using the lmerTest package (Kuznetsova et al. 2017). Bonferroni-adjusted p-values were calculated across PCs and model terms with α = 0.005.

#### Modeling Genome-Wide Allele-Frequency Dynamics with Beta-Binomial GLMMs

To identify SNPs whose allele frequency trajectories differed between A→C and C→A populations, we fit a beta-binomial generalized linear mixed model at each locus. The beta-binomial family accommodates overdispersion inherent to pool-seq allele-count data (Wiberg et al. 2017). At SNP *i*, minor allele counts *Y*_*ijk*_ for population replicate *j* at generation *k*, with *n*_*ijk*_ total reads were modeled as:

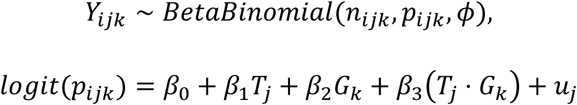

where *p*_*ijk*_ is the underlying allele frequency, *T*_*j*_ is the trajectory indicator (0 = A→C, 1 = C→A), *G*_*k*_ is the continuous generation variable, *β*_1_, *β*_2_, *β*_3_ are fixed-effect coefficient for treatment, generation, and their interaction *u_j_* ∼ *N*(0, *σ_u_*^2^) is the random effect for population replicate *j*, and *ϕ* is the overdispersion parameter. Here the interaction term *β*_3_ captures differential allele frequency change over generations between trajectories, including anti-parallel responses where allele frequencies shift in opposite directions between the A→C and C→A populations.

To correct for multiple comparisons across the genome while accounting for the non-independence of test statistics arising from linkage disequilibrium, we estimated a permutation-based false discovery rate (FDR) following the approach of Jha et al. (2015) as implemented in Kawecki et al. (2021). For each of five permutations, treatment labels were randomly reassigned across the 20 population replicates while preserving each replicate’s complete allele frequency trajectory across timepoints. The beta-binomial GLMM was then refitted at each SNP on the permuted data, generating an empirical null distribution of test statistics under pure drift. The null distribution for FDR estimation was constructed by sorting each permutation’s p-values independently and taking the element-wise minimum across permutations at each rank position. For each model term, the FDR at a given p-value cutoff was estimated as the ratio of the cumulative proportion of null p-values to the cumulative proportion of observed p-values at or below that cutoff, with a monotonicity correction applied to ensure FDR is non-increasing toward more significant p-values. P-values of exactly zero, arising from numerical underflow at loci with extreme allele frequency differentiation, were replaced with the smallest representable positive floating-point value prior to FDR estimation to preserve rank order. SNPs were retained for analysis only if successfully fitted in both the observed data and all five permutations. SNPs were considered statistically significant if they passed both a genome-wide FDR threshold of 0.01 and a minimum effect size of 0.3, applied uniformly to each model term. The 0.3 threshold follows Kawecki et al. (2021) and is intended to exclude SNPs with statistically significant but biologically trivial differences. The effect size metric was defined to reflect the biological hypothesis tested by each term: for the treatment term, the absolute difference in mean allele frequency between A→C and C→A populations averaged across timepoints; for the generation term, the mean absolute allele frequency change over time averaged across trajectories; and for the interaction term, the absolute difference in allele frequency change between trajectories, oriented with respect to the allele at higher frequency in A→C populations at generation 0.

To confirm that significant treatment × generation interactions reflected anti-parallel allele frequency responses, we calculated the change in mean allele frequency between the first and last sampled timepoints for each SNP that met our FDR threshold, averaging across replicates within each trajectory. All frequency changes were expressed relative to the A-allele, defined for each SNP as the allele at higher frequency in A→C populations at generation 0. Where the minor allele frequency in A→C populations at generation 0 exceeded 0.5, the minor allele was treated as the A-allele and frequency changes were sign-flipped accordingly, ensuring that a positive value consistently indicates movement toward the A→C state. The joint distribution of allele frequency changes across the two trajectories was visualized as a hexbin density plot.

Lastly, to facilitate potential future investigation of candidate loci, we identified the most significant marker within independent genomic regions from the set of significant treatment × generation interactions and annotated them with respect to nearby genes. To do this, we divided the genome into non-overlapping 10kb windows along each chromosome, and the SNP with the smallest p-value within each window was recorded as the representative marker for that region. The 100 most significant markers from this list were then annotated using the ‘VariantAnnotation’ Bioconductor package (Obenchain et al. 2014), with transcript annotations from the *Drosophila melanogaster* dm6 reference assembly. For each SNP, we identified all annotated transcripts whose genomic coordinates overlapped the SNP position, including coding exons, introns, three-prime UTRs, five-prime UTRs, splice sites, and the 2 kb promoter window upstream of each transcription start site; SNPs outside all such features were classified as intergenic.

#### Genome-Wide Heterozygosity Analysis

To quantify heterozygosity dynamics across the reciprocal A→C and C→A experimental evolution trajectories, allele frequencies were estimated from pooled sequencing counts at SNPs included in the trajectory dataset. For each sample, major allele counts were obtained by subtracting minor allele counts from total coverage values, and minor allele frequencies were calculated on a per-site basis.

Expected heterozygosity was then calculated independently at each SNP as:

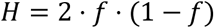

where *f* is the minor allele frequency. Heterozygosity values were averaged across all SNPs for each replicate population at each sampled generation.

Because the trajectory SNP table retains loci polymorphic across the dataset as a whole, sites that appeared fixed within a particular population or timepoint could still contribute to the analysis if variation was present elsewhere in the dataset. Consequently, temporal changes in mean heterozygosity reflect allele frequency dynamics at known polymorphic sites rather than invariant genomic positions. This is the relevant quantity for assessing variation loss and recovery, as monomorphic sites cannot contribute to heterozygosity change regardless of selection regime. Similar approaches have been used in previous experimental evolution studies (Burke et al. 2010; Remolina et al. 2012; Graves et al. 2017).

Changes in mean heterozygosity over time were visualized using boxplots stratified by trajectory. To assess temporal changes in heterozygosity within each trajectory, two-sided Welch’s t-tests were performed comparing replicate-level mean heterozygosity at each post-founding generation to generation 0. Because each trajectory involved a small, ordered family of three comparisons against a common baseline, p-values were adjusted separately within each trajectory using the Holm-Bonferroni sequential correction (Holm 1979). This approach controls the familywise error rate as stringently as a standard Bonferroni correction while gaining power by adapting the rejection threshold to the rank of each p-value. Corrections were applied within trajectory rather than pooled across both trajectories, as A→C and C→A comparisons address distinct directional hypotheses and pooling would impose unnecessary overcorrection. To further investigate the apparent loss of variation observed in the A→C trajectory, we examined whether sites that appeared fixed at generation 0 truly lacked standing variation. This analysis was motivated by the observation that many SNPs in the A→C populations appeared fixed based on standard-coverage sequencing, yet these same populations exhibited rapid phenotypic and genomic rebound when selection was reversed, an outcome inconsistent with widespread true fixation.

To characterize highly differentiated sites in the A→C trajectory, SNP data were obtained from the trajectory SNP table and corresponding deep sequencing counts from the founding A-type1 population. SNPs were identified using the permutation-based FDR framework described above, retaining sites with significant treatment × generation interaction terms (FDR < 0.01, effect size ≥ 0.3). These SNPs were then filtered to identify pseudo-fixed sites in A-type1, defined as positions where the minor allele was undetectable or fixed under standard coverage (minor allele count = 0 or equal to total coverage), and merged with deep sequencing data to compute minor allele frequencies (MAFs), allowing detection of low-frequency alleles that would otherwise be missed.

The distribution of MAFs at pseudo-fixed sites was visualized using histograms and compared against a randomly sampled set of genome-wide SNPs of equal size to provide a null background. To formally test whether pseudo-fixed sites exhibited reduced but nonzero variation, we performed a permutation test in which the mean MAF of pseudo-fixed sites was compared to a null distribution generated by repeatedly sampling random SNP sets from the deep sequencing data (n = 10,000 permutations). Empirical p-values were calculated as the proportion of permuted means less than or equal to the observed mean, and standardized z-scores were used to quantify deviation from the null expectation. Prior to analysis, a minimum minor allele count threshold of 3 was applied to exclude observations potentially attributable to sequencing error. At a median coverage of ∼767x, a conservative Illumina per-base error rate of 0.001 predicts approximately 0.75 spurious reads per site; sampling-with-replacement simulations using a pool of 400 haploid alleles sampled at 767x indicate a minimum detectable frequency of ∼0.00293 (1/341), and the count-3 threshold corresponds to ∼0.00391 (3/767), sitting modestly above this floor. As the deep sequencing was conducted on a pool of 200 females (400 haploid alleles) drawn from a census population of approximately 1,500 individuals, it captures a substantial but incomplete sample of population-level allelic diversity, and sites appearing fixed even at higher depth may retain alleles below this sampling threshold.

To contextualize the role of genetic drift in shaping the low-frequency allele reservoir detected by deep sequencing, we estimated the effective population size (Ne) of each A-type founder population using the poolSeq package (Taus et al. 2017) in R. Allele frequency change was compared between two timepoints separated by approximately 219 generations of continued A-type selection: timepoint 1 of the A-type founder populations as the ancestral sample and timepoint 4 of the same extant populations as the derived sample. Ne was estimated independently for each of the ten matched population pairs using the estimateNe function with the temporal method of Jorde and Ryman (1995), which accounts for variance in allele frequency change attributable to both genetic drift and the additional sampling variance introduced by pool sequencing.

#### Generalized Linear Mixed Model Analysis of ‘newly derived’ versus founder populations

To test the repeatability of genomic evolution under reversed selection, we asked whether populations selected for a given treatment (A or C) converged on the same allele-frequency states as the original founder populations adapted to that regime. Specifically, the ‘founder’ populations in this comparison are the long-standing A-type and C-type progenitor lines that have experienced continued generations of selection under their respective regimes and serve as reference endpoints for genomic convergence. We tested whether C→A populations resemble founder A populations, and whether A→C populations resemble founder C populations. Under repeatable evolution, populations that independently experience the same selection pressure are expected to arrive at similar allele-frequency distributions, despite distinct evolutionary histories.

We fit a quasibinomial generalized linear model at each SNP with population type (newly derived vs. founder) as a fixed effect, following the approach recommended by Wiberg et al. (2017) for allele-frequency comparisons in stratified pool-seq data. We used a quasibinomial GLM here rather than the beta-binomial GLMM applied in the trajectory analysis because this comparison involves only a single timepoint, providing 20 observations per SNP instead of 80. The beta-binomial model’s additional dispersion parameter is better estimated when more observations are available per locus, and at the reduced sample size here it can become unstable. The quasibinomial framework accommodates overdispersion inherent to pool-seq allele counts through an estimated dispersion parameter, which absorbs replicate-level variance without requiring an explicit random effect. Significance of the population-type effect was assessed by t-test on the corresponding coefficient. Chromosome 4 was excluded due to its atypical recombination properties. SNPs showing significant differentiation between newly derived and founder populations were interpreted as evidence of historical contingency or incomplete reversibility, whereas SNPs lacking differentiation were taken as consistent with repeatable convergence on a shared allele-frequency state.

Multiple testing was controlled using the same permutation-based FDR framework described above for the trajectory analysis, with population-type labels (old vs. new) shuffled across the 20 replicate populations. SNPs were considered significantly differentiated if they passed both a genome-wide FDR threshold of 0.01, and we again applied a minimum allele frequency difference of 0.3 between founder and newly derived populations, following Kawecki et al. (2021).

## Data Availability

Raw genomic and phenotypic data are available through Dryad (DOI:10.5061/dryad.tdz08kqdc) and scripts used to process and analyze data are available through Github (https://github.com/krarnold/A-C_Genomic_Trajectory_Project). The DNA sequence data supporting the conclusions of this article are available in the NCBI SRA repository (PRJNA1426013).

## Acknowledgments

This work was supported by the National Institutes of Health Maximizing Investigators’ Research Award to MAP (NIH R35GM155286). We thank the UCI Genomics High-Throughput Facility for their assistance with library preparation, sequencing, and technical support that made this work possible. We thank the developers of Novoalign for providing the alignment software used in processing and mapping our sequencing reads. We also extend our great thanks and appreciation to Dr. Jose Ranz and Dr. Bryan Clifton for their training in DNA extractions and for the use of their laboratory facilities. Additionally, we thank Dr. David Hubert for his valuable feedback on an earlier version of this manuscript. Most importantly, we are grateful to the dozens of undergraduate researchers who assisted with the experiments, without whose help this work could not have been done, especially S. Gagucas, G. Medrano, R. Odina, J. Sue, D. Youn, N. Ly, N. Nguyen, T. Nguyen, E. Choi, K. Hoang-Yanis, C. Banoub, M. Ishizuka, A. Attalla, D. Bruce, C. Wang, E. Brown, B. Khan, J. Hana, A. Vo, K. Co, J. Anup, J. Mai, J. Lee, K. Bryan, R. Chhetri, and B. Ung.

## Author Contributions

M.A.P., M.R.R., and L.D.M. conceptualized and oversaw the project. M.A.P., M.R.R., L.D.M., and K.R.A. designed the experiment. A.P., V.V.C., R.D.R., K.C., C.O.C., and M.Q. collected all fly samples for genomic analyses and carried out mortality assays and larval development experiments under the supervision of K.R.A. K.R.A. extracted samples for genomic sequencing. Z.S.G. and L.D.M analyzed phenotypic data. K.R.A., Z.S.G., and M.A.P. analyzed the genomic data. K.R.A. and M.A.P. wrote the manuscript.

