## Supplemental_Figures.pdf for "Beyond Fixation: Persistent Genetic Variation Under Intense Selection"

**Supplemental Figure Captions**

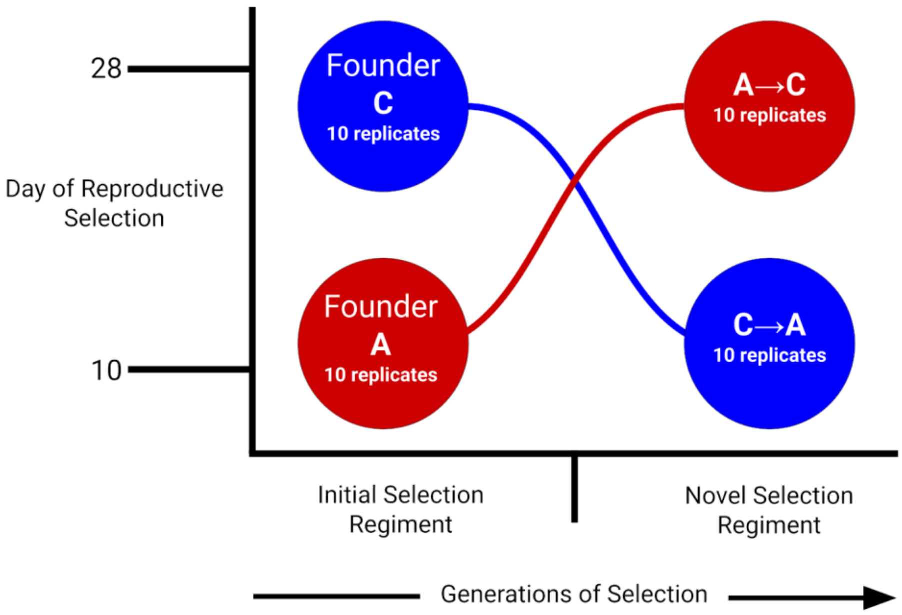


**Figure S1. Experimental design of reciprocal A→C and C→A selection trajectories.** Schematic illustrating the derivation of 20 trajectory populations from 20 long-established founder populations under antiparallel life-history selection. The y-axis represents the day of reproductive selection imposed by each regime, and the x-axis represents generational time. Ten founder C-type populations (blue, day 28) were subjected to A-type selection conditions, generating C→A trajectories selected for progressively earlier reproduction (day 10). Ten founder A-type populations (red, day 10) were subjected to C-type selection conditions, generating A→C trajectories selected for progressively delayed reproduction (day 28). Crossing lines illustrate the antiparallel nature of the two trajectories. Each group consists of 10 independently maintained replicate populations.

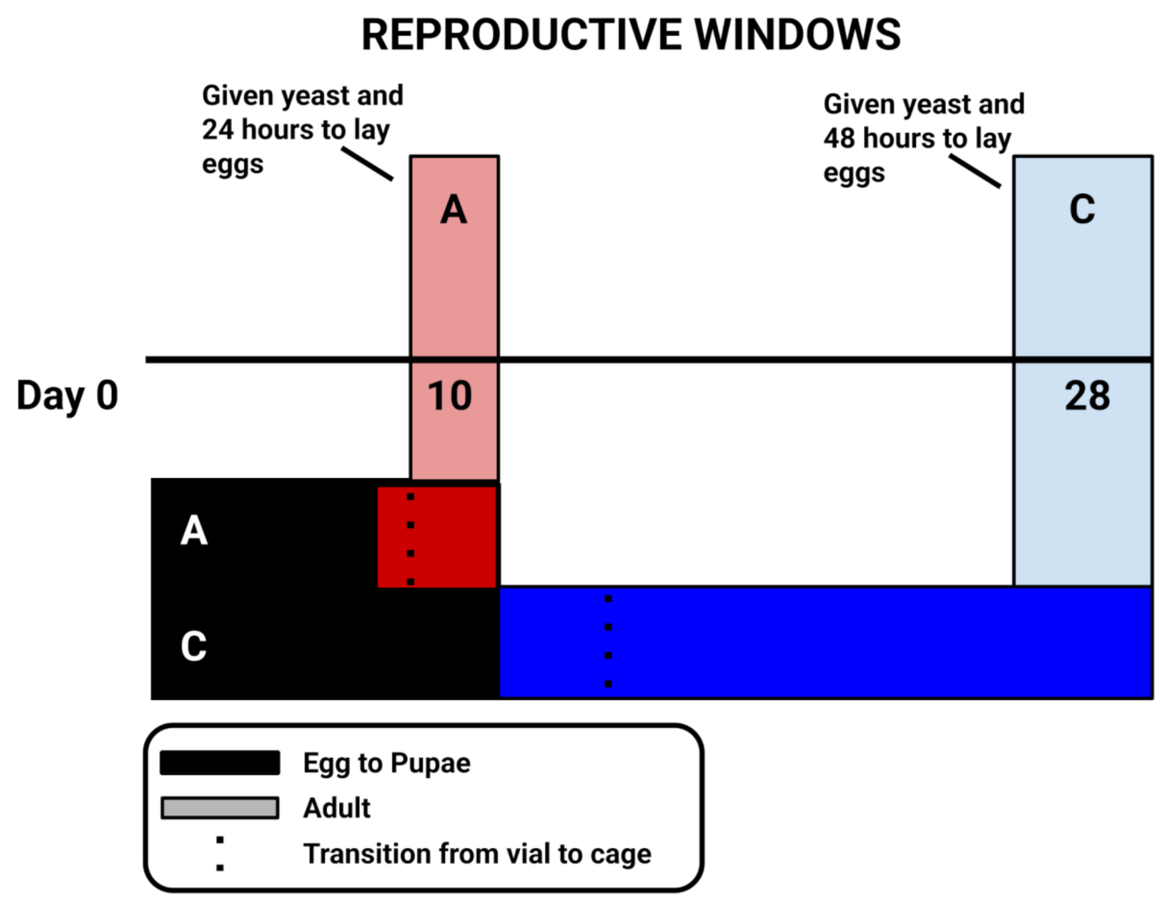


**Figure S2. Schematic Representation of the Life Cycle of Experimental Populations.** The first capitalized letter of a treatment name indicates the type of life-history regime for which it has sustained selection: A (10-day cycle) & C (28-day cycle). The regions in black represent the length of time each treatment spends as pupae before emerging as an adult. The color of the bars denotes the selection regime, A-types represented by red, C-types in blue, and the length of bar marks the adulthood before a new generation is derived. A-type and C-type populations are initially reared in vials for 9 and 14 days, respectively. On day 9 (A-type) or day 14 (C-type), adults are transferred to cages, indicated by the dotted lines, where they are maintained until their designated reproductive windows. Eggs collected during these windows are used to propagate the next generation, maintaining the distinct age-at-reproduction regimes that define each selection.

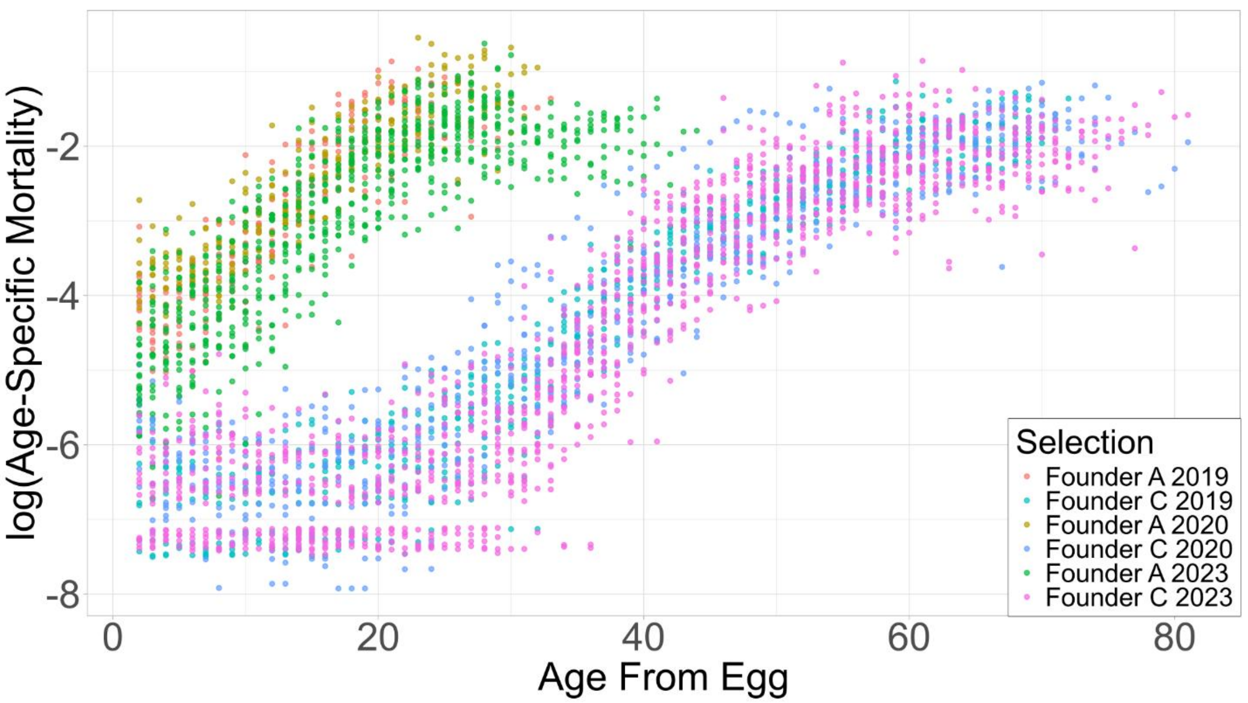


**Figure S3. Founder Age-Specific Mortality Assays, Three Timepoints.** Mortality is expressed as the natural log transformed age-dependent mortality relative to the temporal age from egg for the six mortality distributions. Each dot represents the log mortality rate at a specific age for a given population within each selection regime. Ancestral populations (A&C) served as the controls for every midpoint trajectory mortality assay, making 3 assays for each treatment to assess the year-by-year consistency of the treatments relative to handling effects. Conserved between each experimental year is the pattern of A-types undergoing accelerated death in contrast to the C-types which exhibit a plateau up until Day 28. Significant overlap within selection regimes highlights the reproducibility of the system and relative robustness to any potential handling effects.

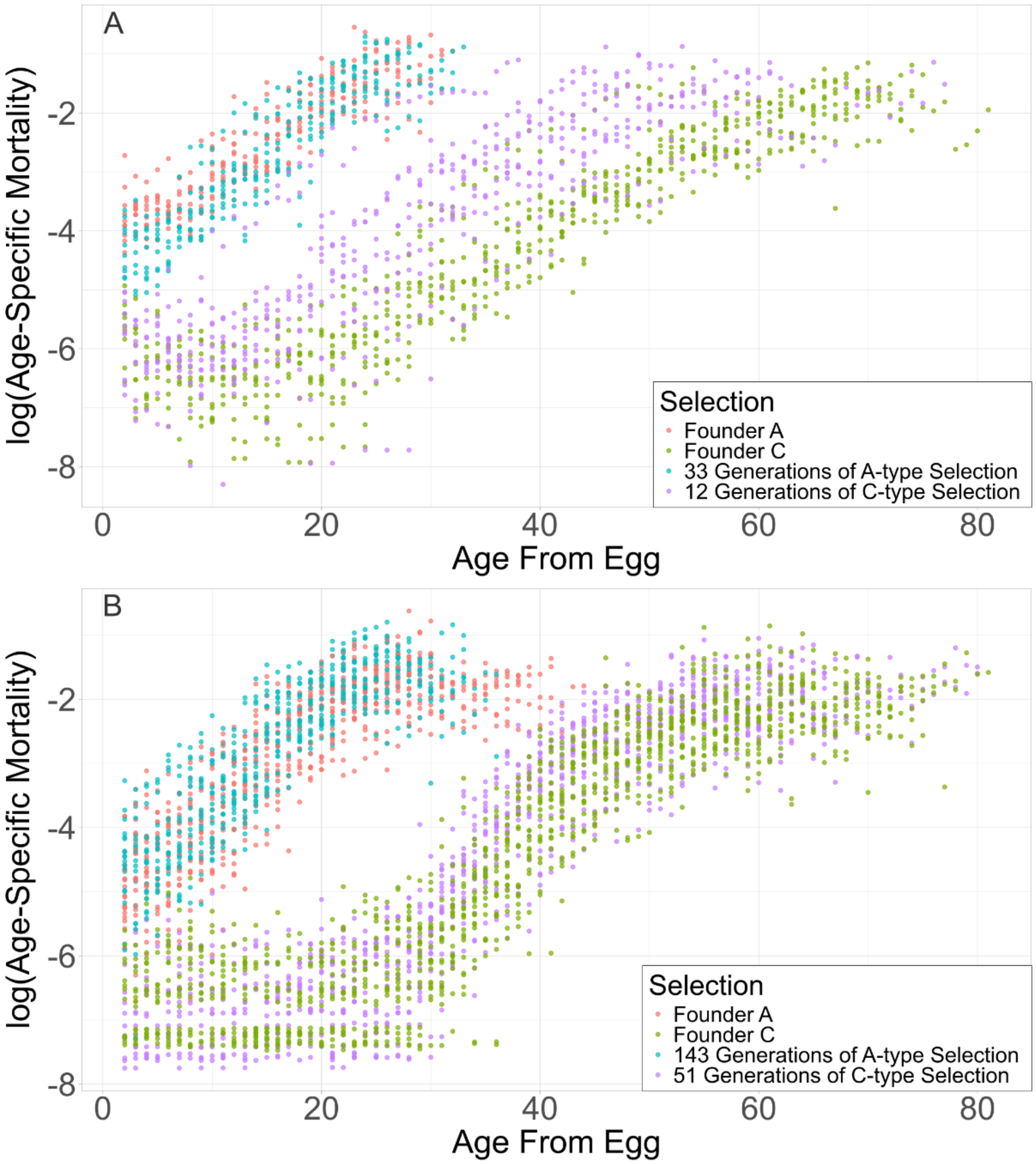


**Figure S4. Age-specific mortality profiles following reciprocal selection.** Age-specific adult mortality plotted as the natural log–transformed mortality rate as a function of age from egg for founder populations and their reciprocally selected derivatives. (A) Early phase mortality assay: A→C populations after ~12 generations of delayed-reproduction selection and C→A populations after ~33 generations of early-reproduction selection. (B) Late phase mortality assay: A→C populations after ~51 generations of delayed-reproduction selection and C→A populations after ~143 generations of early-reproduction selection. Each point represents the mortality rate at a given age for an individual population within each selection regime. Founder A- and C-type populations provide baseline mortality profiles for comparison.

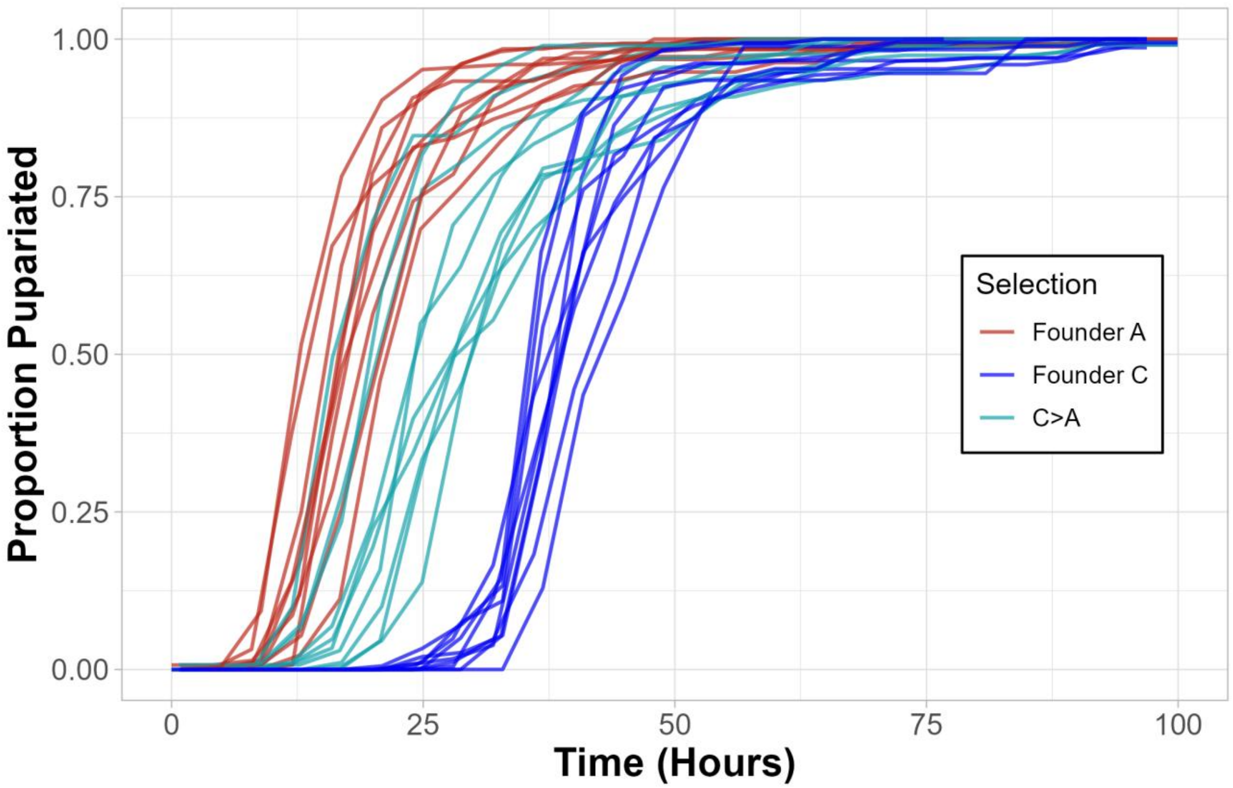


**Figure S5. Cumulative pupariation timing, C→A trajectory, early phase.** Cumulative proportion of individuals that have pupariated, plotted as a function of time from egg (hours) for founder A-type (red), C-type (blue), and derived C→A populations (teal) at approximately 15 generations of early-reproduction selection. Each line represents an individual population replicate. C→A populations show intermediate pupariation timing relative to their respective founders, consistent with partial convergence toward the A-type phenotype at this early assay timepoint.

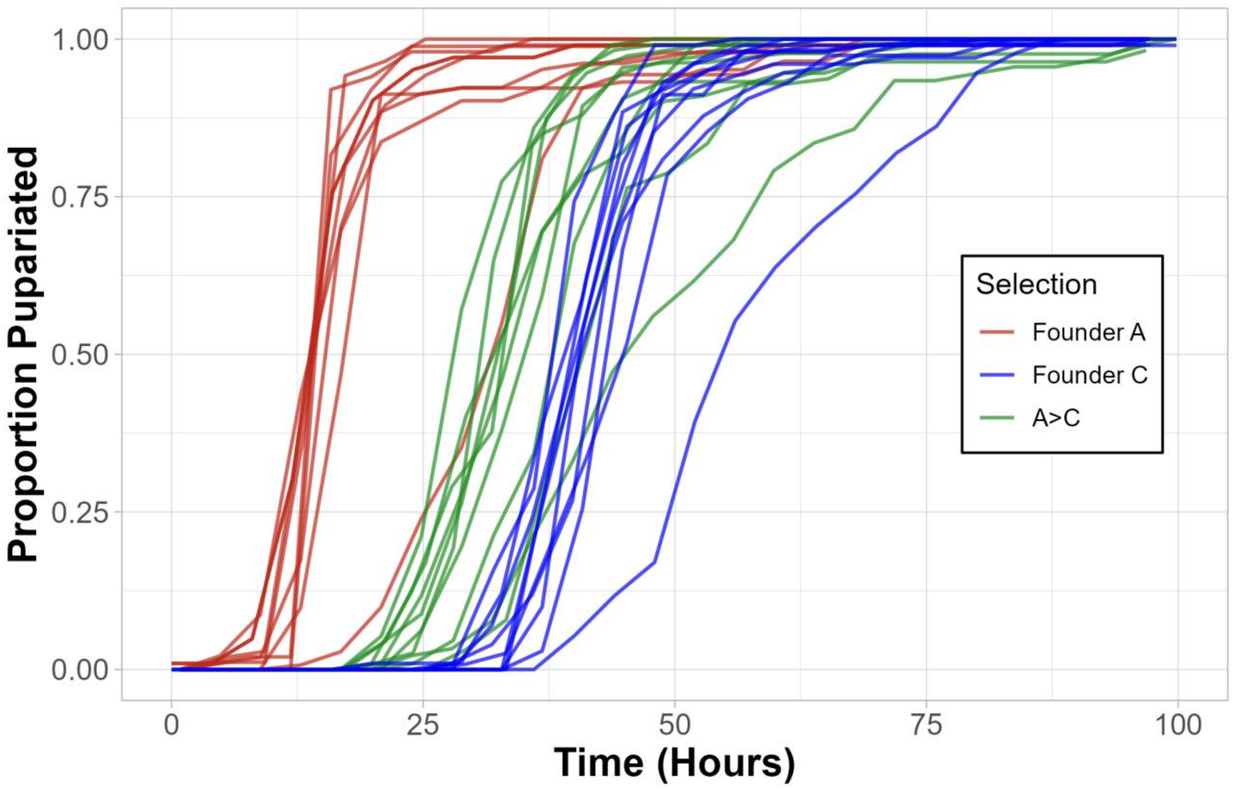


**Figure S6. Cumulative pupariation timing, A→C trajectory, early phase.** Cumulative proportion of individuals that have pupariated, plotted as a function of time from egg (hours) for founder A-type (red), C-type (blue), and derived A→C populations (green) at approximately 6 generations of delayed-reproduction selection. Each line represents an individual population replicate. A→C populations show intermediate pupariation timing relative to their respective founders, consistent with partial convergence toward the C-type phenotype at this early assay timepoint.

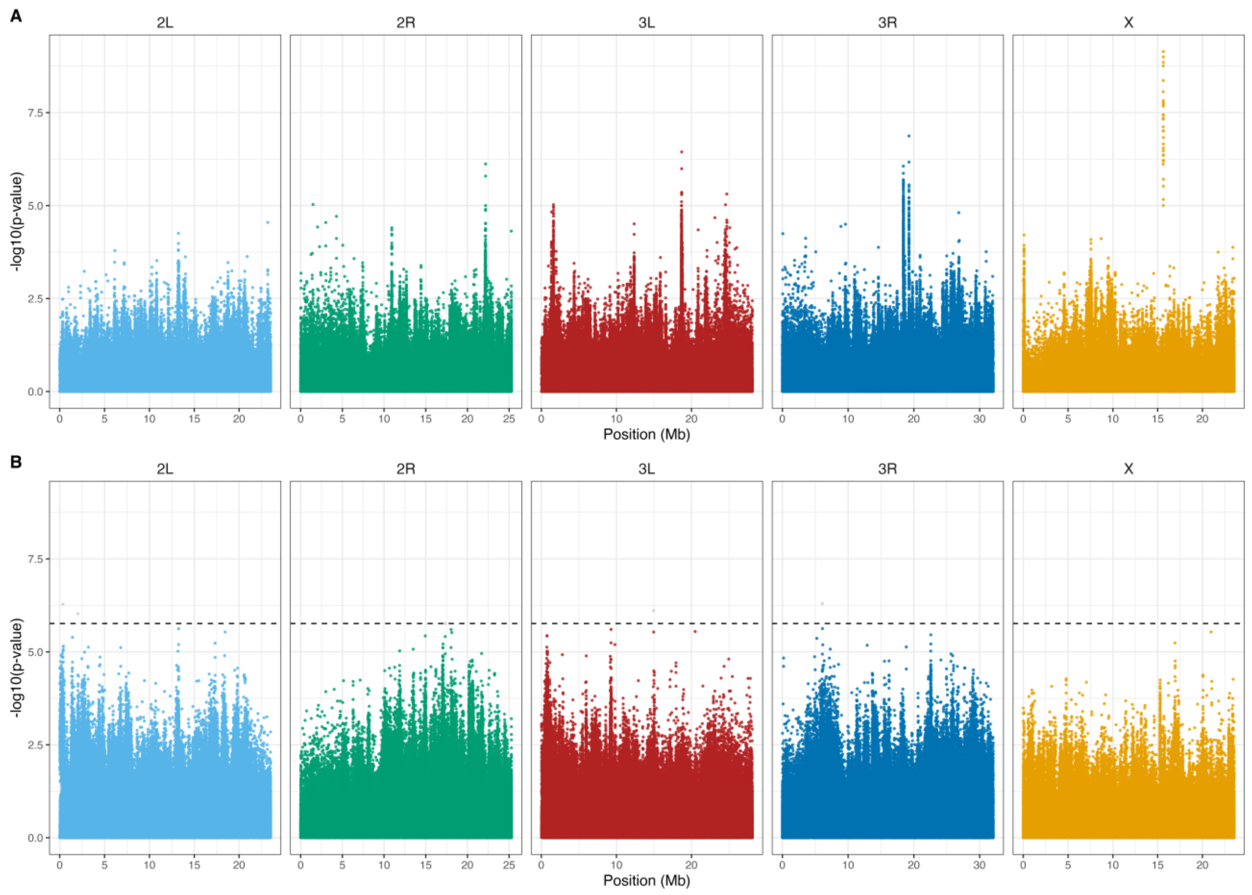


**Figure S7. Genome-wide differentiation between newly derived and long-established founder populations at the final timepoint.** Manhattan plots show quasibinomial GLM results comparing allele frequencies in reciprocally selected populations to long-established founder populations maintained under the same selection regime. Each point represents a SNP plotted by chromosomal position (Mb) on the x-axis and $-\log_{10}$(p-value) on the y-axis. Chromosomes are color-coded: 2L (light blue), 2R (green), 3L (red), 3R (dark blue), and X (gold). (A) A→C populations compared to long-established C-type founder populations after ~65 generations of delayed-reproduction selection. (B) C→A populations compared to long-established A-type founder populations after ~182 generations of early-reproduction selection. The dashed line indicates the permutation-based FDR and effect size threshold (FDR ≤ 0.01, minimum allele frequency difference ≥ 0.3). It was not possible to draw a line for (A) as no SNPs met the FDR threshold. In both cases, no SNP meet the combined FDR and effect size thresholds.
